# Acute high-dose irradiation of human primary ovarian cells reveals a shift in transcriptomics profile and impairment in cell-cell adhesion

**DOI:** 10.1101/2025.03.19.644095

**Authors:** Spyridon Panagiotis Deligiannis, Tianyi Li, Elisabeth Moussaud-Lamodière, Akos Végvári, Anastasios Damdimopoulos, Darja Lavogina, Kiriaki Papaikonomou, Roman Zubarev, Ganesh Acharya, Agne Velthut-Meikas, Pauliina Damdimopoulou, Andres Salumets, Valentina Di Nisio

## Abstract

**STUDY QUESTION:** How do human cortical (cPOCs) and medullary (mPOCs) primary ovarian cells respond to acute X-ray exposure?

**SUMMARY ANSWER:** Acute high-dose X-ray exposure causes a shift in cPOCs and mPOCs transcriptomic profiles and impairs significantly their cell-cell adhesion ability.

**WHAT IS KNOWN ALREADY:** Radiotherapy is a leading cancer treatment, due to its effectiveness in targeting malignant cells. However, it can also affect healthy cells, potentially causing organ dysfunction, among which ovaries. When targeted radiotherapy is not feasible, fertility preservation is recommended to avoid premature ovarian insufficiency. In addition, the effects of irradiation on ovarian somatic cells remain poorly understood.

**STUDY DESIGN, SIZE, DURATION:** Ovarian tissue was obtained from patients undergoing gender-affirming surgery at Karolinska University Hospital Huddinge, Sweden. The ovarian tissue was separated into cortex and medulla, then individually dissociated into single-cell suspensions using mechanical and enzymatic methods. Monolayer cultures from cPOCs and mPOCs were exposed to a single dose of 10 Gy X-ray irradiation or left unexposed as paired controls. Following irradiation, the cells were cultured at various time-points for further molecular and morphological evaluation.

**PARTICIPANTS/MATERIALS, SETTING, METHODS:** Ovarian tissue from 8 patients (age 23-36 years) was used. Dissociated cPOCs and mPOCs were cultured to 80% confluence and irradiated with 10 Gy (1.33 Gy/min), with non-irradiated controls. Cellular ATP and mitochondrial activity were assessed, followed by immunofluorescence staining for canonical irradiation-induced effects in cells: DNA damage, apoptosis and cell cycle progression. Bulk RNA-sequencing was performed on controls and irradiated samples. Libraries were prepared using the Illumina Stranded mRNA Prep Ligation protocol and sequenced on Illumina NovaSeq6000 platform. Genes were considered to be differentially expressed under the cut-off of false discovery rate (FDR) < 0.05. Subsequently, affected biological pathway was predicted using all expressed genes ranked by log_2_ fold change again hallmark gene sets. To further investigate the potential upstream regulators, transcription factor enrichment analysis were performed based on DEGs. To assess changes at protein level, we mapped the proteomic profile using liquid chromatography-tandem mass spectrometry. Peptides were considered to be differentially expressed (DEPs) under the cut-off of p-value < 0.01. Finally, we measured the ability of cPOCs and mPOCs to form 3D aggregates after seeding irradiated and non-irradiated cells on Biosilk scaffolds.

**MAIN RESULTS AND THE ROLE OF CHANCE:** Following irradiation, ATP levels and mitochondrial activity in cPOCs and mPOCs were comparable to controls, indicating minimal irradiation impact on cell viability and proliferation. Immunofluorescence analysis confirmed the modulation of canonical pathways, such as DNA damage, apoptosis and cell cycle in both cPOCs and mPOCs. Transcriptomic analysis showed that cPOCs and mPOCs at 1 h post-irradiation clustered together with the related 1 h control. However, a shift in transcriptomic profile was observed after 4 h and even more after 24 h post-irradiation in both cPOCs and mPOCs. Gene set enrichment analysis (GSEA) indicated upregulation of the p53 pathway at 4 h and 24 h post-irradiation, alongside downregulation of MYC targets, E2F targets, the G2/M checkpoint and mTORC1 pathway. Gene pattern analysis showed irradiation-dependent trends related to extracellular matrix (ECM) organisation, p53-mediated apoptotic mechanisms and chromosome segregation during the 24 h period following irradiation. Additionally, transcription factor enrichment analysis based on DEGs suggested p53 and MYC as potential upstream regulators. On a proteomic level, DEPs associated with ECM organisation and cytoskeleton formation were detected at 4 and 24 h post-irradiation. Finally, X-ray exposure hindered the cell-cell adhesion ability of both cPOCs and mPOCs, leading to impaired formation of Silk-Ovarioids.

**LARGE SCALE DATA:** The RNA sequencing count matrix is deposited in Gene Expression Omnibus (GEO) with accession number GSE291604. The mass spectrometry proteomics data have been deposited to the ProteomeXchange Consortium via the PRIDE partner repository with the dataset identifier PXD061796. The code used for the analysis can be found at https://github.com/tialiv/X-Ovary.

**LIMITATIONS, REASONS FOR CAUTION:** The ovarian tissue was obtained from gender-affirming surgery patients who received androgen treatment before removal. Even though unlikely, this hormonal treatment might influence ovarian environment and alter cellular response to irradiation. Additionally, in this study the impact of X-ray exposure was assessed on a monolayer cell model, thus limiting the extrapolation power of our results to ovary *in vivo*. Lastly, the impact of irradiation was focused on the somatic cell populations that are essential for ovarian function. Further studies are needed to investigate the effects of X-ray exposure on ovarian follicles and their function.

**WIDER IMPLICATIONS OF THE FINDINGS:** Understanding the roles of MYC, p53 and cell adhesion factors in response to irradiation could guide the development of future ovarian protective strategies. These findings lay the foundation for further studies on ovarian tissue protection and fertility preservation in cancer patients.

**STUDY FUNDING/COMPETING INTEREST(S):** This work was funded by the European Union’s HORIZON 2020 research and innovation programme (MATER) under the Marie Skłodowska-Curie Actions (grant agreement No: 813707), the Estonian Research Council (grants PRG1076 and PSG608), the Orion Research Foundation sr personal grant, the Research grant from the Center for Innovative Medicine (CIMED) and the Karolinska Institutet Consolidator Grant.

## Introduction

The rising global incidence of cancer, affecting nearly 9 million women in 2022 (Cao et al., 2024), highlights the urgent need for advancements in cancer treatment. Ionizing irradiation or radiotherapy (RT), often used in combination with chemotherapy and surgery, is a cornerstone of cancer therapy, and it is applied to approximately 30-50% of patients (Barton et al., 2014). The widespread use of RT is attributed to its low overall cost – accounting for only 5% of total cancer care expenditures - and its significant contribution to cancer survival, responsible for approximately 16% of patients being cured (Bentzen et al., 2005).

While RT is effective in targeting cancer cells, it also impacts normal non-malignant cells, increasing the risk of long-term collateral damage and compromising tissue integrity. This has been linked to adverse effects in various organs (Fernström et al., 2024; Lawrie et al., 2018; Pouvreau et al., 2024). Ovarian tissue is among the affected organs, sensitive on RT-induced side effects (Shin et al., 2020).

Ovarian tissue consists of two distinct compartments: the outer cortex and the inner medulla. The cortex, about 1 mm thick, contains small pre-antral follicles including the dormant ones constituting the ovarian reserve. The medulla has a smoother texture and houses larger secondary and antral follicles. Single cell RNA sequencing revealed different profile of cell populations in medulla compared to cortex, especially with a high diversity in granulosa cells (GC) (Fan et al., 2019; Wagner et al., 2020), possibly due to multiple sub-populations of GC from antral follicles (Roos et al., 2022). Together, these layers support important ovarian functions such as folliculogenesis and steroidogenesis.

Despite the cellular complexity of the ovarian tissue, most studies reported irradiation-related effects on follicles. In early-stage follicles, granulosa cells exhibit high proliferative activity, while the oocyte is highly sensitive to stressors and toxicants. This vulnerability makes early-stage follicles particularly susceptible to radiation damage, potentially leading to premature ovarian insufficiency (POI) and infertility (Immediata et al., 2022; Kimler et al., 2018; Mishra et al., 2017; Wang et al., 2023). The extent of damage depends on different factors such as age of the patient, radiation dose, and irradiation field (Meirow and Nugent, 2001). For instance, a single dose of 2 Gray (Gy) irradiation in pelvic area can damage over 50% of the ovarian reserve, while doses of 10 Gy may cause cessation of ovarian function (Irtan et al., 2013). Clinically, doses of 10 – 20 Gy in children and 4 – 6 Gy in adults are associated with POI (Chemaitilly et al., 2006). In 2021, based on a stringent systematic literature review, the PanCareLIFE Consortium published guidelines for recommendation of fertility preservation (*i.e.,* oocyte cryopreservation, embryo cryopreservation, ovarian tissue cryopreservation) in female patients with childhood, adolescent and young adult cancer. The guidelines strongly recommend counselling the patients about options for fertility preservation in cases of RT with ovary in the field (Mulder et al., 2021). However, the impact of RT is not only confined to follicles. In fact, it can also significantly affect the ovarian stroma, which have been observed to shrink following irradiation due to fibrosis and atrophy. Moreover, RT can cause small vessel damage leading to thrombosis and creation of fibrin masses in the lumen, thus further compromising ovarian functions (Stroud et al., 2009).

In this study, we investigated the effects of ionizing radiation on human primary ovarian cells (POCs) *in vitro*. To address this, POCs were isolated from the ovarian cortex (cPOCs) and medulla (mPOCs) and exposed to a single 10 Gy dose of X-ray irradiation. We then assessed cellular ATP levels and mitochondrial reductase activity over a 7-day period, interrogated the activation of canonical pathway related to irradiation-induced damage, analysed the changes in cellular transcriptomic and proteomic profiles, and evaluated the ability of ovarian cells to form 3D ovarian aggregates following X-ray irradiation.

## Material and Methods

### Study setup

In this study, we investigated the impact of X-ray irradiation on primary ovarian cells (cPOCs and mPOCs), obtained from gender-affirming patients (n=8). We performed a screening for the selection of the most relevant dose (2, 4, and 10 Gy) and recovery timings (4, 24, 48, 72, 120, 168 h post-irradiation) using a granulosa cell-like tumor cell line (KGN) (n=3). A dose of 10 Gy was applied to cPOCs and mPOCs, and intracellular ATP levels (n=8) and mitochondrial activity (n=8) were assessed at different time points over a 7-day culture period post-irradiation. Subsequently, we evaluated through immunofluorescence analysis (n=5) the activation of well-known canonical pathways altered after irradiation at three time points (1, 4, 24 h post-irradiation), namely DNA damage, apoptosis and cell cycle. Furthermore, transcriptomic (n=5) and proteomics analysis (n=5) was performed (1, 4 and 24 h post-irradiation). Finally, the functionality of irradiated cPOCs and mPOCs was tested by evaluating their ability to form 3D ovarian aggregates *in vitro* (n=2).

### Study participants and ethical approval

Human ovarian tissue samples were obtained from patients undergoing gender-affirming surgery (n=8; age range: 23-36 years) at Karolinska University Hospital Huddinge, Sweden. Signed written informed consent was obtained from all participants in accordance with the Declaration of Helsinki, permitting the use of their ovarian tissue for research purposes.

During surgery, bilateral oophorectomy was performed to remove ovarian tissue, part of which was immediately transferred to the laboratory within 10 min in Dulbecco’s Phosphate Buffered Saline (DPBS) supplemented with calcium, magnesium, glucose and sodium pyruvate (Life Technology, UK). The ovarian medulla was carefully trimmed and separated from the cortex (depth 1-1.5 mm), and the samples were cryopreserved using a slow-freezing protocol as previously described (Schmidt et al., 2003). Briefly, cortical and medullary tissues were cut into small pieces of 5 x 5 x 1 mm in size and equilibrated on ice for a maximum of 30 min in a slow-freezing medium containing 7.5% ethylene glycol (Sigma-Aldrich), 33.9 mg/ml sucrose (Sigma-Aldrich) and 10 mg/ml human serum albumin (HSA, Vitrolife) diluted in DBPS (Thermo, Scientific) on a shaker. Following incubation, the tissue pieces were transferred into Nunc^TM^ CryoTube^TM^ Vials (ThermoFisher Scientific) containing 1 ml of slow-freezing medium and placed into a controlled-rate freezer (Kryo 230-1.7, Planer PLC). The tissues were subsequently thawed and used in later experiments.

To ensure confidentiality, all samples were pseudonymised using randomised codes for subsequent experiments and data analysis. Personal information was managed in compliance with the European General Data Protection Regulation (GDPR).

This study was approved by the Swedish Ethical Review Authority (reference number 2024-08606-01).

### KGN cell culture

KGN cells were chosen for dose optimisation due to their resemblance in proliferation rates and transcriptomics profile to POCs (Tarvainen et al., 2023). KGN was obtained from RIKEN Cell Bank (RBRC-RCB1154, Japan). Cell was verified by Eurofins Genomics Europe Applied Genomics GmbH using short tandem repeat analysis to exclude the possibility of contamination.

KGN cells were cultured in DMEM/F12 (Life Technology, USA) supplemented with 10% heat-inactivated fetal bovine serum (HI-FBS; Life Technology, USA), 1% penicillin-streptomycin (Life Technology, USA) and 1% 100X Glutamax^TM^ (Life Technology, USA). Cells were maintained at 37°C in a humidified atmosphere containing 5% CO_2_.

For experiments, cells were seeded in 96 well plates, depending on the assay further performed. Cell densities are reported in Supplementary Table 1. Media was refreshed every two days. Cells were maintained at 37 °C in a humidified atmosphere with 5% CO_2_, with media changes every other day.

### Ovarian tissue dissociation and primary ovarian cell culture

For tissue dissociation, the cryopreserved/thawed ovarian cortex and medulla were independently digested into a single cell suspension using a mechanical and enzymatic method, as previously described (Wagner et al., 2020). Briefly, tissue pieces were finely cut into small pieces using scalpels, and further enzymatically dissociated with 1 mg/ml collagenase IA (Sigma Aldrich, USA), 50 μg/ml Liberase^TM^ (Roche Diagnostics, Germany), and 10 IU/ml DNase I (Roche Diagnostics, Germany) diluted in Dulbecco’s Modified Eagle Medium F12 (DMEM/F12; Life Technology, USA) supplemented with 2.5% HI-FBS (Life Technology, USA). Digestion was carried out in a 37 °C shaking water bath for 40-50 minutes.

Enzymatic dissociation was terminated by adding equal volume of DMEM/F12 (Life Technology, USA) supplemented with 10% HI-FBS (Life Technology, USA), followed by a 5 min centrifugation at 300x g (centrifuge 5702R; Eppendorf) to collect POCs. Cells were resuspended in Dulbecco’s Modified Eagle Medium, low D-glucose (1 g/L), pyruvate (DMEM low glucose, pyruvate; Gibco, Life Technology, USA), supplemented with 10% HI-FBS (Life Technology, USA), and 1% penicillin-streptomycin (Life Technology, USA).

The cell suspension was filtered through a 40 μm cell strainer (VWR, USA), and cells with diameter between 7.5-25 μm were counted using the Moxi mini automated cell counter (OR-FLO Technologies). Isolated cPOCs and mPOCs were seeded in different densities as reported in Supplementary Table S1, depending on the experimental design. Cultures were maintained at 37 °C in a humidified atmosphere with 5% CO_2_, with media changes every other day.

### KGN and POCs X-ray irradiation

KGN, cPOCs and mPOCs were subjected to X-ray irradiation using the Xstrahl CIX2 system (Xstrahl), operated at 195 kV and 10 mA, delivering a dose range of 1.33 Gy/min. Irradiation was conducted in X-ray Irradiation Core facility at Karolinska Institutet (Huddinge, Sweden). A 3 mm aluminium filter was applied during exposure, and the cells were rotated to ensure uniform dose distribution, with a Focus-to-Skin Distance (FSD) of 40 cm.

To identify the optimal irradiation dose for the primary cell studies, KGN cells were exposed to total doses of 2 Gy, 4 Gy, or 10 Gy. The maximum dose of 10 Gy was selected based on the PanCareLIFE Consortium guidelines (Mulder et al., 2021). Additionally, 2 Gy and 4 Gy doses were included, as 2 Gy has been shown to damage over 50% of the ovarian reserve (Irtan et al., 2013; Wallace et al., 2003), and 4 Gy has been associated with POI in adults (Chemaitilly et al., 2006).

Following irradiation at these doses, KGN cells were used for the evaluation of cellular ATP levels and mitochondrial activity. Based on the results of this screening, a total dose of 10 Gy was selected for subsequent irradiation experiments involving cPOCs and mPOCs.

### Measurement of intracellular ATP level and mitochondrial activity

CellTiter-Glo® assay (Promega, USA) was used to assess cell viability of KGN (n=3), cPOCs (n=8) and mPOCs (n=8) within 7 days of culture following irradiation based on the intracellular ATP levels, following manufacturer’s instructions. Briefly, equal volume of CellTiter-Glo® reagent and medium was mixed on an orbital shaker to induce cell lysis, followed by 10 min incubation at room temperature in the dark to stabilise the luminescent signal. After incubation, the luminescence intensity was read using a SpectraMax i3x ELISA plate reader (Molecular Devices, Germany) at a 578 nm wavelength.

For the quantification of irradiation-induced damage, the changes in metabolic activity were measured in KGN (n=3), cPOCs (n=8) and mPOCs (n=8) within 7 days of culture following irradiation using the Cell Proliferation Kit I (MTT) (Roche Diagnostics, Germany), according to the manufacturer’s instructions. Briefly, cells were mixed with MTT labelling reagent followed by 4 h incubation in a humidified atmosphere and an overnight incubation with Solubilization buffer. The absorbance was then measured in SpectraMax i3x ELISA plate reader (Molecular Devices) at a 550 nm wavelength and reference wavelength >650 nm.

### Immunofluorescence staining

Irradiated cPOCs (n=3) and mPCOs (n=3) were fixed with 4% paraformaldehyde (PFA; Sigma-Aldrich) at 1 h, 4 h and 24 h post-irradiation for immunofluorescence staining. As controls, non-irradiated cPOCs and mPOCs were cultured and fixed concomitantly with the 1 h irradiated samples. Cells were permeabilised with blocking buffer containing DPBS (Life Technologies, Paisley, UK), 0.3% Triton X-100 (Sigma-Aldrich), 3% normal donkey serum (NDS; Nordic Biosite) and 5% bovine serum albumin (BSA; Sigma-Aldrich). Subsequently, cells were incubated overnight at 4 °C on an orbital shaker with primary antibody (Suppl. Table S2) diluted in antibody buffer containing DPBS (Life Technologies, Paisley, UK) and 1% NDS (Nordic Biosite). Negative controls were incubated with blocking buffer instead of the primary antibody.

After primary antibody incubation, cells were washed in washing buffer (PBST) containing DPBS (Life Technologies, Paisley, UK) and 0.3% Triton X-100 (Sigma-Aldrich) and subsequently incubated for 1 h with secondary antibody (Suppl. Table S2) diluted in antibody buffer. When primary antibodies from the same host species were used for different markers in the same section, a double-blocking procedure was performed to prevent cross-reactivity. Briefly, at the end of the first secondary incubation process, slides underwent a second round of blocking before the overnight incubation with the same-host primary antibody. 4’,6-diamidino-2-phenylindole dihydrochloride (DAPI; ThermoFischer Scientific) was used for nuclear counterstain (1:1000 dilution in antibody buffer). Slides were mounted using DAKO Fluorescence Mounting Medium (Agilent Technologies, USA).

Cells were imaged using a 20x air objective with 1.5x lens on a confocal microscope (Nikon Eclipse Ti2) with channels detected with emission filters accordingly: 438/24 blue, 511/20 green, 560/25 red, and 685/40 far-red, with a Kinetix sCMOS camera (Crest Optics). Figures were processed in Omero figure software, contrast and brightness were adjusted according to the signal expression and kept uniform in each panel.

We used CellProfiler (version 4.2.5) to create image analysis pipelines for immunofluorescence image quantification (Stirling et al., 2021). Briefly, we corrected the images for non-homogeneous illumination. Then, we created a mask to delimit single-cells suspensions as regions of interest (ROIs) based on Gaussian-filtered and dilated DAPI signals from the illumination-corrected images. We identified individual cells based on the nuclei fluorescence signals. The fluorescence intensity of each marker from the whole cells was then measured individually and compiled using RStudio. The fluorescence intensity was visualised as ratio for protein expression.

### RNA extraction and library preparation

RNA extraction was performed using the RNeasy Micro Kit (Qiagen, Germany) following the manufacturer’s instructions. Briefly, cells were lysed in 75 μl of Buffer RLT (Qiagen, Germany). To eliminate contamination of genomic DNA, DNase I (Qiagen, Germany) treatment was applied. RNA was eluted with 14 μl of RNase-free water and the concentration was measured using a NanoPhotometer (IMPLEN, Nordic Biolabs). RNA quality was further assessed through the Agilent Bioanalyzer 2,000 (Agilent, USA). Only samples with RNA integrity value (RIN) > 9 and A260/A280 > 1.8 were used for library construction. Libraries were prepared using the Illumina Stranded mRNA Prep Ligation protocol (Illumina, USA) according to the manufacturer’s instruction, with 10 ng of RNA used as input.

### RNA sequencing and data analysis

RNA sequencing (paired-ends, 2 × 150 bp) was conducted on the Illumina NovaSeq6000 platform at the Bioinformatics and Expression Analysis (BEA) core facility, Karolinska Institute, Sweden. *Fastq* files were trimmed using TrimGalore (version 0.6.1) to remove the low-quality reads before mapping. STAR aligner (version 2.7.11a) was used to map the trimmed *fastq* files to human genome hg38. Subsequently, the bam files were aligned using *featureCount* function in subread package (version 2.0.3) and annotated using UCSC gtf file (available at https://emea.support.illumina.com/sequencing/sequencing_software/igenome.html). The pipeline used for trimming, mapping, and alignment was provided in a snakemake file (snakemake version 5.10.0). Differential expression analysis was performed in R (version 4.3.2) through Rstudio (R Core Team, 2022) using the DESeq2 package. Comparisons of 24 h control vs 1 h control, 1 h irradiated vs 1 h control, 4 h irradiated vs 1 h control, 24 h irradiated vs 24 h control were performed separately in cortex and medulla dataset. Differentially expressed genes (DEGs) with false discovery rate (FDR) < 0.05 were considered statistically significant. Gene pattern analysis was performed on DEGs identified in all the comparisons in cortex and medulla separately using DEGreport package (Pantano, 2024). Subsequently, the signalings related to the identified gene clusters were inferred using Gene Ontology (GO) over-representation analysis. Affected biological pathways were identified using Gene set enrichment analysis (GSEA) against the MsigDB (Liberzon et al., 2011) hallmark gene set on all expressed genes ranked by log_2_ fold change through clusterProfiler (Wu et al., 2021; Yu et al., 2012), pathview (Luo and Brouwer, 2013), DOSE (Yu et al., 2015) and apeglm (Zhu et al., 2019) packages. ComplexHeatmap (Gu, 2022; Gu et al., 2016) package was used for heatmap plotting. Codes used for analysis are available at https://github.com/tialiv/X-Ovary.

For transcription factor (TF) enrichment we obtained the links between the TFs and their targets from TFLink (https://tflink.net) and restricted it to TFs that had been detected by chromatin immunoprecipitation. To assess whether a TF is enriched in significantly regulated genes we used hypergeometric testing in R.

### Proteomics sample preparation

Irradiated cPOCs (n=5) and mPOCs (n=5), along with their respective controls, were snap-frozen and thawed on ice and lysed by adding 10 µL of 8 M urea, followed by sonication in a water bath for 5 min. Next, 70 µL of 0.5 M NaCl in 50 mM Tris-HCl (pH 8.5) and 0.8 µL 100x of protease inhibitor (Roche Diagnostic, Germany) were added. After an additional 5 min of sonication, the lysates were centrifuged at 13,000 *g* for 10 min at 4 °C. Protein concentration was measured using BCA assay (Pierce, USA). A volume of lysate corresponding to 7.7 µg of protein was supplemented with 1 M urea and 438 mM NaCl in Tris-HCl buffer, bringing the final volume to 75 µL. Proteins were reduced by adding 2.8 µL of 250 mM dithiothreitol (Sigma-Aldrich, USA) and incubated at 37 °C for 45 min while shaking at 400 rpm on a block heater. For alkylation, 3.1 µL of 500 mM iodoacetamide (Sigma-Aldrich, USA) was added, and the samples were incubated at room temparature for 30 min while shacking at 400 rpm in the dark. Proteolytic digestion was carried out by adding 0.4 µg sequencing grade modified trypsin (Promega, USA) and incubated for 16 h at 37°C. The digestion was terminated by adding 4.5 µL of concentrated formic acid (FA) and incubating the solutions at room temperature for 5 min. The sample was cleaned using a C18 Hypersep plate with 40 µL bed volume (Thermo Fisher Scientific, USA) and dried in a vacuum concentrator (Eppendorf, Germany).

Samples were labeled with TMT-10plex (Thermo Fisher Scientific, USA) isobaric reagents. Peptides were solubilized in 70 µL of 50 mM triethylammonium bicarbonate and mixed with 100 µg TMT-10plex reagents in anhydrous acetonitrile (ACN) and incubated at room temperature for 2 h. The unreacted reagents were quenched with 6 µL of hydroxyamine at room temperature for 15 min. Biological samples were then combined, dried in vacuum and cleaned on C18 Hypersep plate.

### Liquid Chromatography-Tandem Mass Spectrometry Data Acquisition

The TMT-10plex labeled peptide samples were reconstituted in solvent A, and approximately 2 µg of the samples were injected onto a 50 cm long EASY-Spray C18 column (Thermo Fisher Scientific) connected to an Ultimate 3000 nanoUPLC system (Thermo Fisher Scientific). A 90-minute gradient was used: 4-26% solvent B (98% ACN, 0.1% FA) for 90 min, 26-95% for 5 min, and 95% solvent B for 5 min, with a flow rate of 300 nL/min. Mass spectra were acquired using a Q Exactive HF hybrid quadrupole-Orbitrap mass spectrometer (Thermo Fisher Scientific), scanning from *m/z* 375 to 1700 at a resolution of R = 120,000 (at *m/z* 200), targeting 1×10^6^ ions for a maximum injection time of 80 ms. This was followed by data-dependent higher-energy collisional dissociation (HCD) fragmentation of the top 18 precursor ions with charge states from 2+ to 7+, using a 45-s dynamic exclusion. Tandem mass spectra were acquired with a resolution of R = 60,000, targeting 2×10^5^ ions for a maximum injection time of 54 ms, with quadrupole isolation width set to 1.4 Th and normalized collision energy set to 34%.

### Proteomics data analysis

The raw data files obtained were processed using Proteome Discoverer v3.0 (Thermo Fisher Scientific) with the MS Amanda v2.0 search engine, referencing the human protein database (SwissProt, 20,022 entries, downloaded on February 9, 2023). Up to two missed cleavage sites were allowed for full tryptic digestion, with precursor and fragment ion mass tolerances set to 10 ppm and 0.02 Da, respectively. Carbamidomethylation of cysteine was considered a fixed modification, while oxidation of methionine, deamidation of asparagine and glutamine, and TMT6plex (+229.163 Da) on lysine and peptide N-termini were treated as dynamic modifications. The initial search results were filtered to a 5% FDR using the Percolator node in Proteome Discoverer. Quantification was based on the intensities of the reporter ions.

Proteomics data was analysed using R (version 4.3.2) and RStudio. After removing missing values, data was normalized to median, removed batch effects (through *removeBatchEffect* function in limma (Ritchie et al., 2015) package), and log_2_-transformed before differential expression analysis. Fold changes were calculated by subtracting the mean of the irradiated groups from the mean of the control groups. Comparisons of 1 h irradiated vs 1 h control, 4 h irradiated vs 1 h control, and 24 h irradiated vs 24 h control were performed separately in cPOCs and mPOCs. Subsequently, p-values were calculated using t-test. Peptides were considered to be differentially expressed if the p-values were lower than 0.01.

### Formation and culture of 3D ovarian aggregates

Monolayer cultures of, cPOCs (n=2) and mPOCs (n=2) were detached using TryplE (ThermoFisher Scientific) 24 h post-irradiation and utilised for Silk-Ovarioids formation, as previously described (Di Nisio et al., 2024). Briefly, 20 μl drops of Biosilk^TM^ were pipetted into 24-well plate and air bubbles were introduced into each drop to create dense foams. Irradiated and non-irradiated (control) cPOCs and mPOCs were seeded onto the foams at a density of 1.2 x 10^5^ cells/foam. The seeded foams were incubated at 37 °C for 20 min to allow stabilisation. After stabilisation, each foam was overlaid with 700 μl of culture media and incubated at 37 °C in a humidified atmosphere of 5% CO_2_ for 14 days, with daily media changes.

On day 14, the seeded foams were carefully detached using a spatula, divided into two equal portions, and transferred into a flat-bottom ultra-low attachment 24-well plate (Corning, USA) for further free-floating culture. Silk-Ovarioids were cultured for an additional 40 days with daily media changes. At the end of the culture period, the Silk-Ovarioids were fixed in 4% methanol-free formaldehyde for subsequent morphological evaluation.

### Statistical Analysis

Data from CellTiter-Glo® and MTT assays were processed using GraphPad prism version 10. The Shapiro-Wilk normality test was performed using the dplyr R package to evaluate data distribution. Statistical significance for parametric data was assessed using one-way ANOVA, followed by the Dunnett’s multiple comparisons test. For non-parametric data, statistical significance was then determined using the Kruskal-Wallis test, followed by the Benjamini-Hochberg post-hoc test from the dunn.test R package.

## Results

### Study setup

This study aimed to investigate the effects of X-ray irradiation on primary ovarian cells. After a dose-finding study in KGN cells, a dose of 10 Gy was applied to cPOCs and mPOCs, and intracellular ATP levels and mitochondrial activity were assessed over a 7-day culture period post-irradiation. Transcriptomic and proteomic analyses were conducted at 1 h, 4 h and 24 h post-irradiation with non-irradiated controls sampled at 1 h and 24 h. Immunofluorescence staining was performed in cPOCs and mPOCs at 1 h, 4 h and 24 h post-irradiation, and in non-irradiated 1 h controls. Finally, the functionality of irradiated cPOCs and mPOCs was tested by evaluating their ability to form 3D ovarian aggregates (Silk-Ovarioids) (Fig. 1).

**Figure 1.**
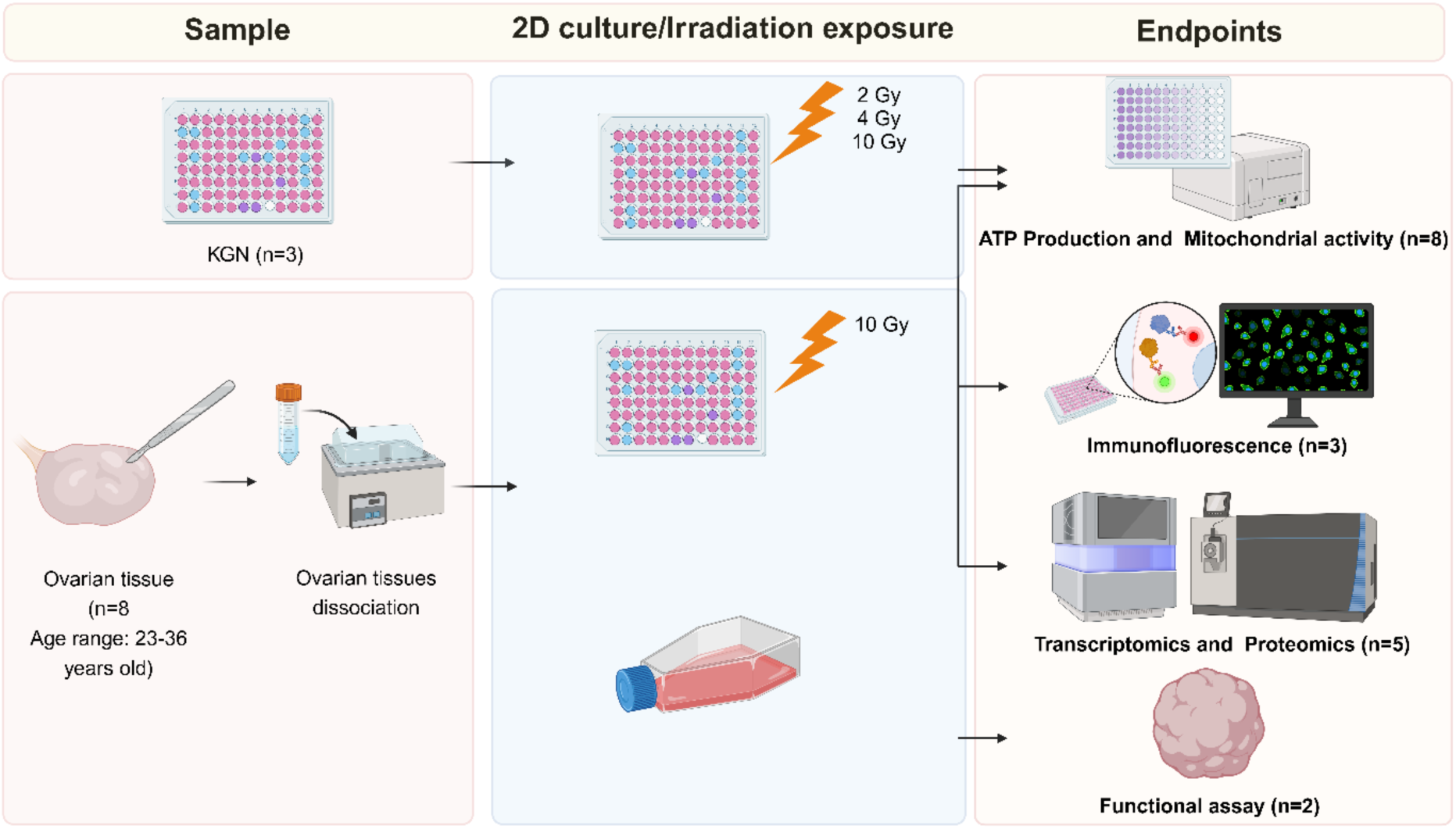
Schematic overview of the experimental setup. Experimental setup of 2D and 3D *in vitro* culture of ovarian primary cells, X-ray exposure and endpoints of the study. ATP, adenosine triphosphate; cPOCs, cortex-derived primary ovarian cells; mPOCs, medulla-derived primary ovarian cells; Gy, Gray. Created with BioRender.com.

### Intracellular ATP levels and mitochondrial activity of KGN and POCs are not significantly affected by irradiation

To identify a relevant and sublethal dose of irradiation, KGN cells (n=3) were exposed to 2 Gy, 4 Gy and 10 Gy of X-ray, and intracellular ATP levels and mitochondrial activity were measured at 4 h, 24 h, 48 h, 72 h, 120 h, and 168 h post-irradiation. No significant differences across all doses and time points were detected (Supp. Fig. 1A-F).

Based on these results, the 10 Gy dose was selected for later experiments involving cPOCs and mPOCs (n=8). ATP levels and mitochondrial activity were monitored at 4 h, 24 h, 72 h, 120 h, and 168 h post-exposure, with unexposed cells at 4 h and 120 h serving as controls. In both cPOCs and mPOCs, ATP levels remained stable throughout the culture period. (Suppl. Fig. S1A, B). Mitochondrial activity showed no significant changes observed across the 168 h post-irradiation culture period (Suppl. Fig. S2C, D).

### Immunofluorescence of canonical pathways induced by irradiation

Since no significant changes were observed in ATP production and mitochondrial activity, we performed immunofluorescence staining on canonical pathways related to irradiation-induced damage (*i.e.,* DNA damage, apoptosis and cell cycle) in cPOCs and mPOCs at 1 h, 4 h and 24 h post-irradiation.

#### Irradiation effect on DNA damage in cPOCs and mPOCs

We selected the well-known DNA damage markers phosphorylated Chk1 (Ser345) (p-Chk1) and γ-H2AX (Ser139) for immunolabeling in both cPOCs and mPOCs (Fig. 2). In cPOCs, protein expression of both p-Chk1 and γ-H2AX peaked at 1 h, and steadily decreased at 4 h, until reaching again levels comparable to control baseline at 24 h post-irradiation (Fig. 2B-C). On the other hand, mPOCs showed a different pattern of response to X-ray exposure. Specifically, p-Chk1 and γ-H2AX proteins reached the maximum levels at 4 h, to then return to baseline levels at 24 h post-irradiation, indicating a transient activation of DNA damage response (Fig. 2E-F).

**Figure 2.**
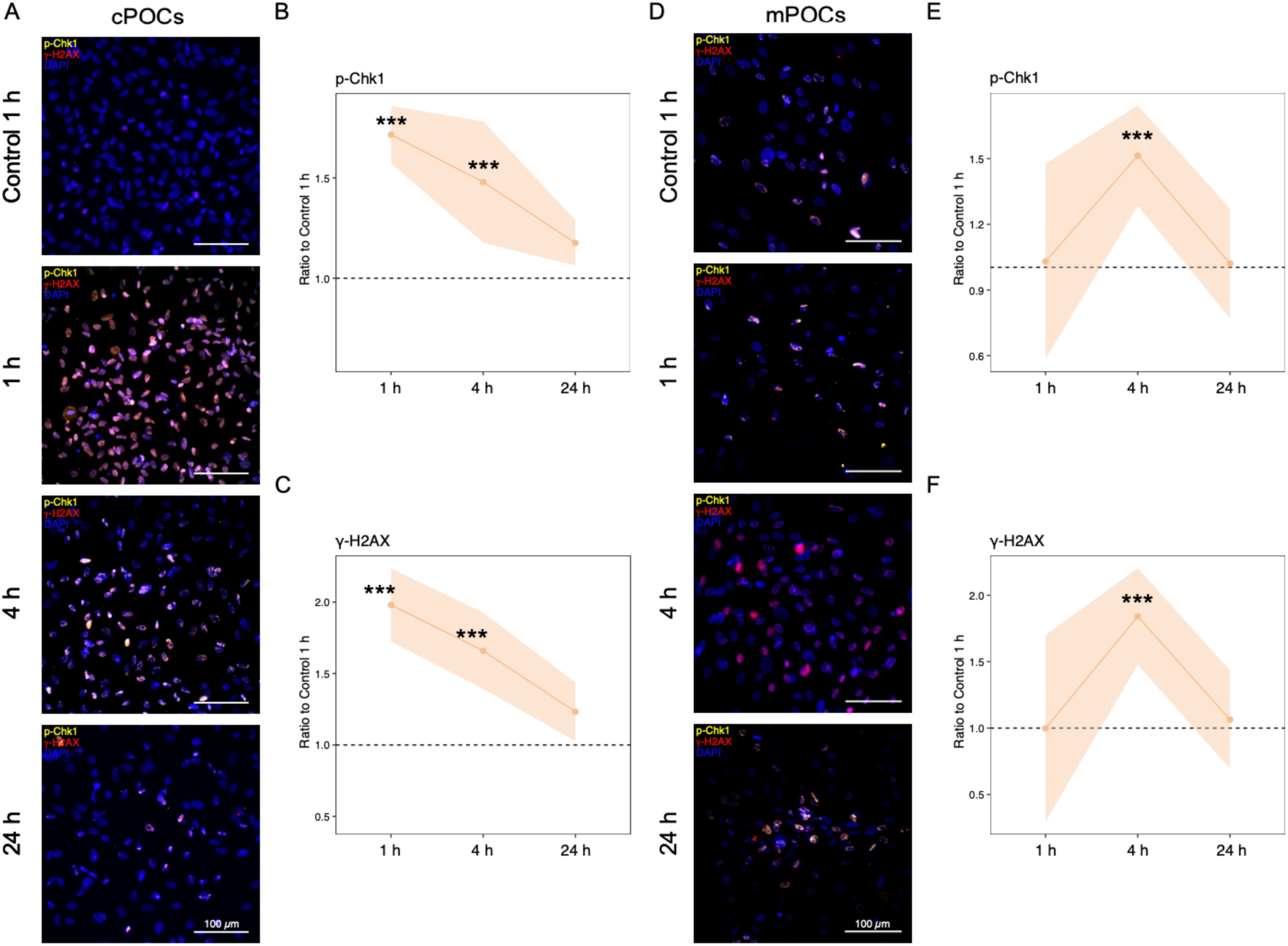
Expression of DNA damage related markers in cPOCs and mPOCs before and after irradiation. Representative images of **(A)** cPOCs and **(D)** mPOCs immunolabeled with DNA damage-related markers: γ-H2AX (red) and p-Chk1 (yellow). Scale bar represents 100 μm. Ratio of irradiated samples to 1 h control at different time points for **(B, E)** p-Chk1 and **(C, F)** γ-H2AX. The mean fluorescent intensity value is represented in yellow. The shaded area around the dots demonstrates the standard deviation. Ratio for protein expression was calculated by dividing the mean fluorescent intensity of each irradiated sample by the mean fluorescent intensity. The statistical significance was analysed using Kruskal-Wallis test with Dunn’s correction and displayed in asterisks (*). *** p > 0.001.

#### Irradiation effect on activation of apoptosis in cPOCs and mPOCs

We assessed the activation of apoptotic mechanisms through immunostaining of cleaved caspase 3 (cl-Cas 3), phosphorylated p53 (Ser15) (p-p53) and Bcl-2 (Fig. 3). In cPOCs, the protein levels of pro-apoptotic markers p-p53 and cleaved caspase 3 showed a significant decrease at 24 h post-irradiation compared to the baseline control (Fig. 3B-C). Contrarily, Bcl-2 was significantly upregulated at the early time points (Fig. 3D). Similarly, in mPOCs the expression of phosphorylated protein of pro-apoptotic markers followed a mirroring trend compared to cPOCs (Fig. 3F-G), as well as the anti-apoptotic marker Bcl-2 (Fig. 3H).

**Figure 3.**
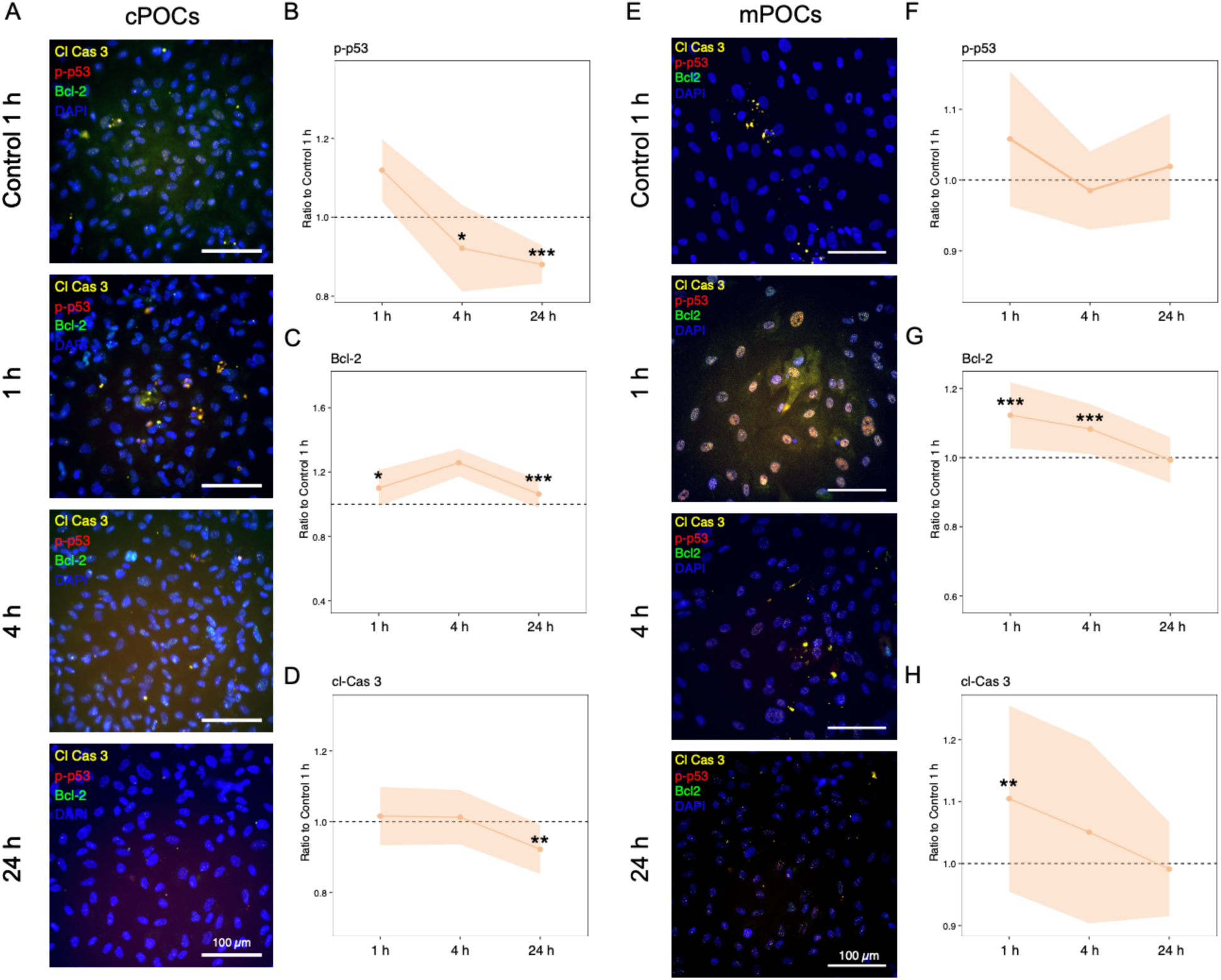
Expression of apoptosis-related markers in cPOCs and mPOCs before and after irradiation. Representative images of **(A)** cPOCs and **(E)** mPOCs immunolabeled with apoptosis-related markers p-p53 (red), Bcl-2 (green), and cl-Cas 3 (yellow). Scale bar represents 100 μm. Ratio of irradiated samples to 1 h control at different time points for **(B, F)** cl-Cas 3, **(C, G)** p-p53, and **(D, H)** Bcl-2. The mean fluorescent intensity value is represented in yellow. The shaded area around the dots demonstrates the standard deviation. Ratio for protein expression was calculated by dividing the mean fluorescent intensity of each irradiated samples by the mean fluorescent intensity The statistical significance was analysed using Kruskal-Wallis test with Dunn’s correction and displayed in asterisks (*).* p < 0.05; ** p > 0.01; *** p > 0.001.

#### Irradiation effect on cell cycle in cPOCs and mPOCs

Finally, we assessed markers related to cell cycle, *i.e.* cyclin D1, cyclin E and phosphorylated p21 (Ser146) (p-p21) in both cPOCs and mPOCs (Fig. 4). In cPOCs, the protein level of both cyclin D1 and E displayed similar trends, with cyclin E being statistically significant across time points (Fig. 4B-C), while the p-p21 expression remained stable (Fig. 4D). In mPOCs, a similar trend was observed in cyclin D1 and E proteins with a significant downregulation at 24 h post-irradiation (Fig. 4F-G). Mirroring cPOCs, p-p21 was significantly upregulated throughout the tested time points (Fig. 4H).

**Figure 4.**
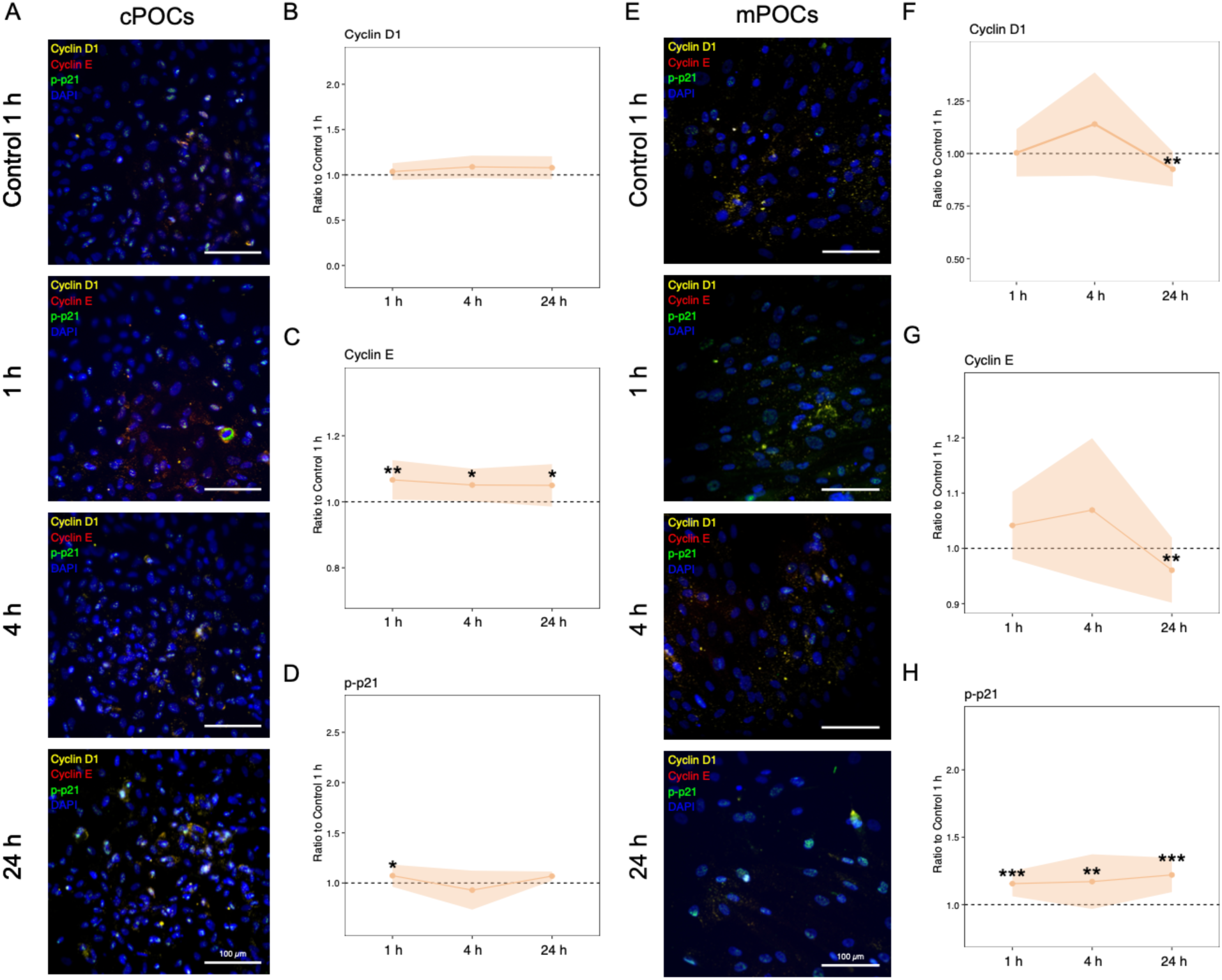
Expression of cell cycle-related markers in cPOCs and mPOCs before and after irradiation. Representative images of **(A)** cPOCs and **(E)** mPOCs immunolabeled with cell cycle-related markers cyclin D1 (yellow), cyclin E (red), and p-p21 (green). Scale bar represents 100 μm. Ratio of irradiated samples to 1 h control at different time points for **(B, F)** cyclin D1, **(C, G)** cyclin E and **(D, H)** p-p21. The mean fluorescent intensity value is represented in yellow. The shaded area around the dots demonstrates the standard deviation. Ratio for protein expression was calculated by dividing the mean fluorescent intensity of each irradiated sample by the mean fluorescent intensity. The statistical significance was analysed using Kruskal-Wallis test with Dunn’s correction and displayed in asterisks (*). p < 0.05; * p < 0.05; ** p > 0.01; *** p > 0.001.

### Transcriptomic analysis identified distinct gene expression profiles between two controls in cPOCs and mPOCS

To assess the impact of X-ray irradiation on transcriptomic profiles, bulk RNA-sequencing was performed on cPOCs and mPOCs (n=5) at 1 h, 4 h and 24 h post-irradiation. Non-irradiated controls were included at 1 h and 24 h. Principal components analysis (PCA) revealed distinct transcriptomic profiles between the 1 h and 24 h controls, in cPOCs (Suppl. Fig. S3A-B) and mPOCs (Suppl. Fig. S3D-E), respectively. Comparison of 24 h and 1 h controls revealed 4,510 DEGs in cPOCs and 2,091 DEGs in mPOCs (FDR < 0.05, Suppl. Table S3). Gene Set Enrichment Analysis (GSEA) revealed upregulation of E2F targets and G2M checkpoint pathways in the 24 h control cPOCs and mPOCs compared to the 1 h control (Suppl. Table S5, Suppl. Fig. S3C, F), indicating that the recorded transcriptomic changes were mainly due to proliferation and cell-cycle progression. Therefore, the early time-points (1 h and 4 h) were compared to the non-irradiated control at 1 h, while the longer time-point (24 h) was compared to the non-irradiated control at 24 h.

### Transcriptomic profile of cPOCs and mPOCs is altered after irradiation

At 1 h post-irradiation, minimal transcriptomic changes were observed (3 DEGs in cPOCs and 1 in mPOCs, Suppl. Table S4). However, significant changes emerged at 4 h with 2,810 and 2,540 DEGs in cPOCs and mPOCs, respectively. These transcriptomic alterations persisted at 24 h with 1,796 DEGs in cPOCs and 2,802 in mPOCs (Suppl. Table S4). In general, PCA across all groups revealed that PC1 separated early time points (1 h and 4 h) from late time points (24 h), while PC2 distinguished 24 h irradiated samples from their respective controls, indicating that the effects of irradiation became more prominent at 24 h (Fig. 5A, B, Suppl. Fig. S4A, B).

**Figure 5.**
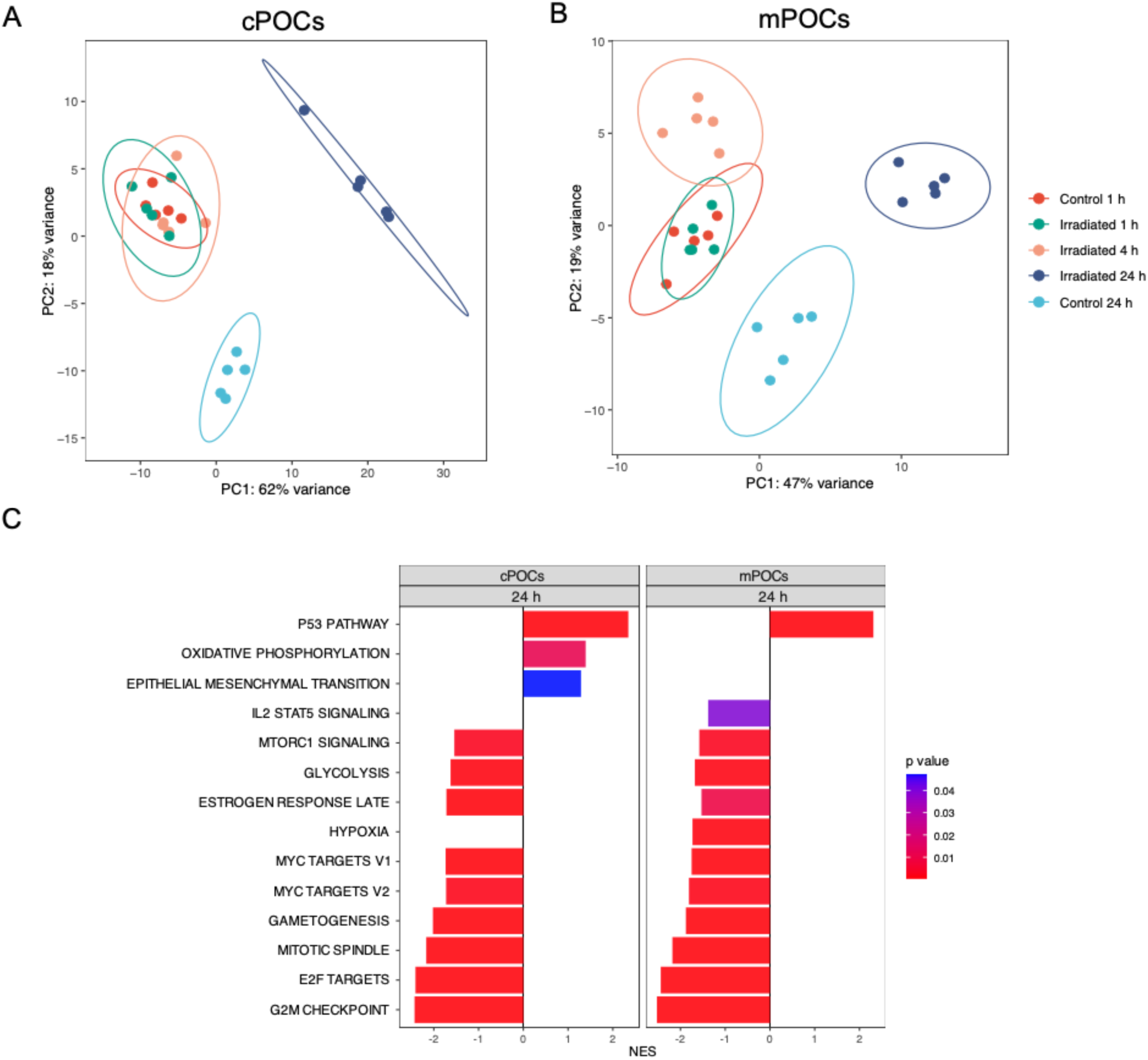
Irradiation induced changes in the transcriptomic profiles of cPOCs and mPOCs. Principal component analysis (PCA) of the top 500 highly variable genes after removing batch effects in the **(A)** cortex and **(B)** medulla. **(C)** Significantly enriched hallmark gene sets predicted by GSEA using all expressed genes ranked by log_2_ fold changes at 24 h post-irradiation. Normalized enrichment score (NES) is shown on the x axis. cPOCs, cortex-delivered primary ovarian cells; mPOCs, medulla-delivered primary ovarian cells.

To assess the biological impact of X-ray irradiation, we performed Gene Set Enrichment Analysis (GSEA) against the hallmark gene sets using all expressed genes ranked by log_2_ fold change. At earlier timepoints, our results showed a significant upregulation of MYC targets, E2F targets and G2M checkpoint pathways at 1 h in cPOCs and at 4 h in mPOCs (Suppl. Fig. S5). Moreover, an upregulation of p53 pathway was observed at 4 h post-irradiation in both cPOCs and mPOCs (Suppl. Fig. S5). The transcriptomic changes became more prominent at 24 h post-irradiation. Overall, the affected pathways after irradiation were relatively similar in cPOCs and mPOCs at 24 h. At the latest timepoint, molecular signatures revealed downregulation of MYC targets, mTORC signalling, E2F targets, G2M checkpoints and glycolysis, and upregulation of p53 pathway (Fig. 5C, Suppl. Table S6). Additionally, we observed some distinct responses, such as upregulation of oxidative phosphorylation and epithelial-mesenchymal transition in cPOCs, and downregulation of hypoxia in mPOCs after 24 h from the X-ray exposure (Fig. 5C, Suppl. Table S6). Due to the more prominent transcriptomic changes 24 h post-irradiation, the subsequent studies focussed mainly on the 24 h effects post-irradiation.

### Gene pattern analysis and GO enrichment revealed effects on the extracellular matrix, DNA damage, and chromosome segregation in cPOCs post-irradiation

To assess whether X-ray irradiation could provoke consistent changes over time, from 1 h to 24 h post-irradiation, we analysed DEGs from all the timepoint comparisons and identified the main gene patterns and their related Gene Ontology (GO) terms. To focus our attention on the most representative irradiation-related gene patterns, we set a maximum of 6 distinct gene clusters in both cPOCs and mPOCs (Fig. 6 and 7). In cPOCs, the 307 genes grouped in cluster 2 exhibited a downregulation at 4 h post-irradiation followed by a sharp upregulation at 24 h above the control baseline (Fig. 6A). As shown by the GO analysis, this pattern was associated with extracellular matrix (ECM) organisation (Fig. 6B and Suppl. Table S7). Moreover, the 378 genes grouped in cluster 5 showed a steady upregulation following X-ray irradiation up to 24 h, whereas the 24 h control remained similar to the 1 h control (Fig. 6A). GO terms enriched in this cluster were vesicle organisation and apoptotic pathways linked to both p53 and DNA damage (Fig. 6C and Suppl. Table S7). Lastly, the 556 genes grouped in cluster 6 displayed a sudden downregulation at 24 h post-irradiation (Fig. 6A). This pattern was related to nuclear chromosome segregation, sister chromatid division and microtubule cytoskeleton organisation (Fig. 6D and Suppl. Table S7).

**Figure 6.**
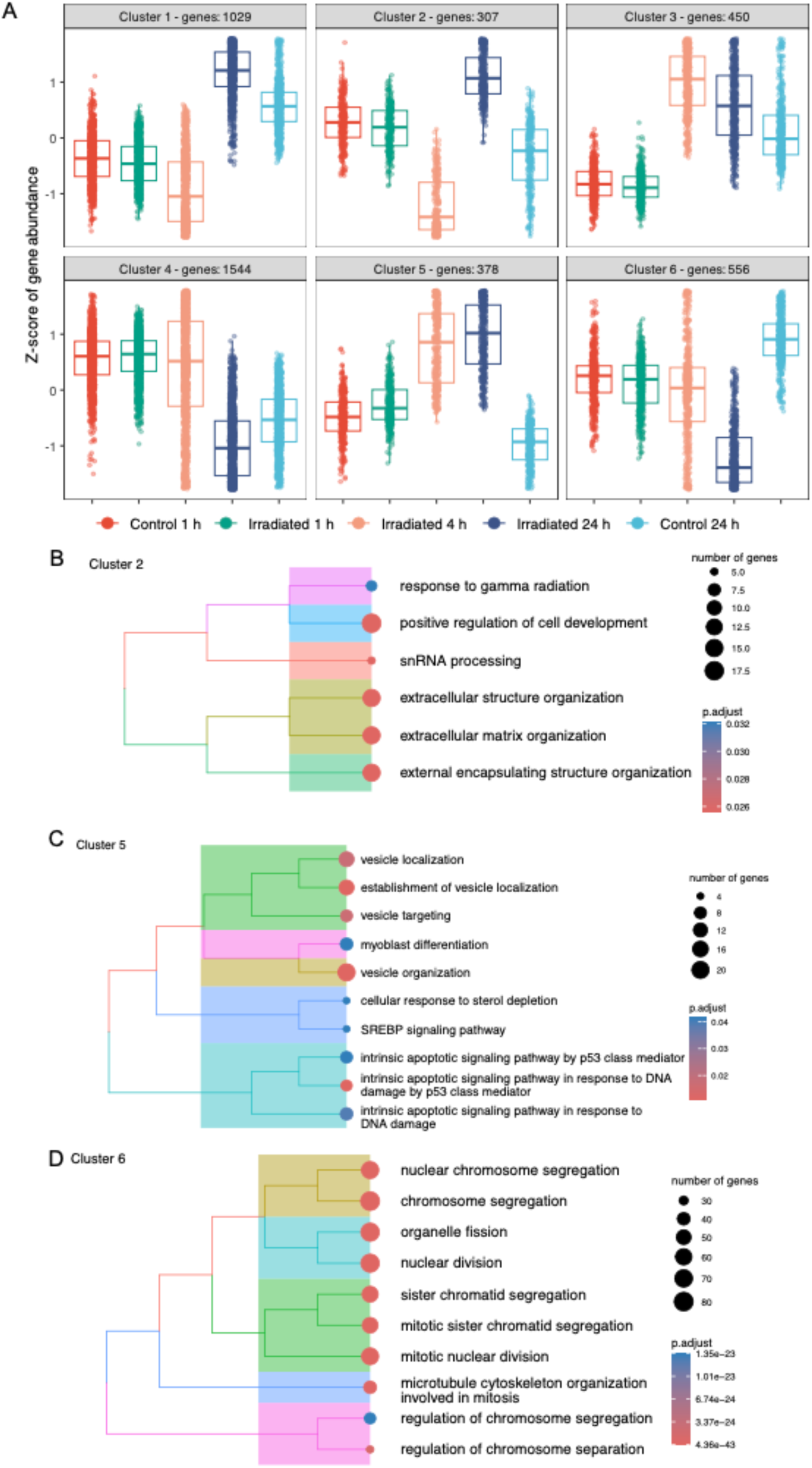
Gene pattern analysis in cPOCs. **(A)** Gene pattern analysis using differentially expressed genes (DEGs) identified in 1 h irradiated cPOCs *vs* 1 h control, 4 h irradiated cPOCs *vs* 1 h control and 24 h irradiated cPOCs *vs* 24 h control. Z-scores of gene abundance is displayed on y-axis. Treeplot of top 10 significantly enriched gene ontology (GO) terms using DEGs identified in **(B)** cluster 2, **(C)** cluster 5 and **(D)** cluster 6. GO terms were clustered using ward.D method based on the similarity of genes in the pathways and the position of the pathway in the GO network. Cluster-enriched terms were shown next to the GO pathways. The size of the dots represents the number of genes enriched in the GOs. The color of the dots indicates the adjusted p-value.

**Figure 7.**
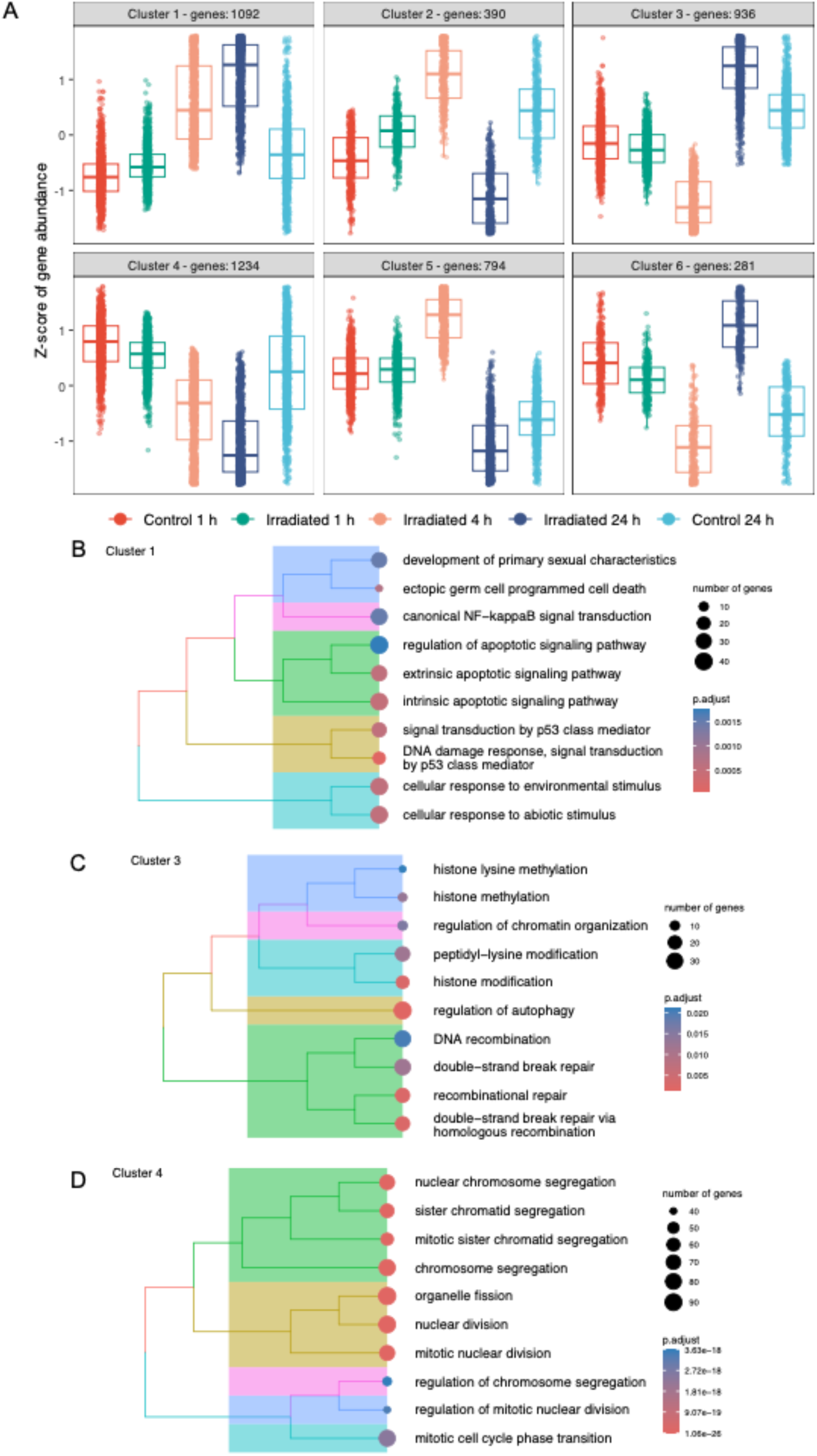
Gene pattern analysis in mPOCs. **(A)** Gene pattern analysis using differentially expressed genes (DEGs) identified in 1 h irradiated mPOCs *vs* 1 h control, 4 h irradiated mPOCs *vs* 1 h control, and 24 h irradiated mPOCs *vs* 24 h control. Z-scores of gene abundance is displayed on y-axis. Treeplot of top 10 significantly enriched gene ontology (GO) terms using DEGs identified in **(B)** cluster 2, **(C)** cluster 5, and **(D)** cluster 6. GO terms were clustered using ward.D method based on the similarity of genes in the pathways and the position of the pathway in the GO network. Cluster-enriched terms are shown next to the GO pathways. The size of the dots represents the number of genes enriched in the GOs. The color of the dots indicates the adjusted p-value.

### Gene pattern and GO analysis revealed effects on DNA damage, histone modification and chromosome segregation in mPOCs

In mPOCs, the 1,092 genes grouped in cluster 1 showed a steep and sharp upregulation up to 24 h post-irradiation, compared to the respective controls (Fig. 7A). Mirroring cluster 5 in cPOCs, GO analysis showed that this cluster was associated to cell death, apoptotic pathway, p53 and DNA damage (Fig. 7B and Suppl. Table S8). The 936 genes grouped in cluster 3 exhibited a drop in expression at 4 h post-irradiation followed by a sharp increase after 24 h, compared to the respective controls (Fig. 7A). This unique signature was connected to double strand break repair, histone lysine modification and regulation of autophagy (Fig. 7C and Suppl. Table S8). Lastly, the 1,234 genes grouped in cluster 4 underwent a constant downregulation during the culture up to 24 h post-irradiation, whereas the 24 h control remained at the same level as the 1 h control (Fig. 7A). Similar to cluster 6 in cPOCs, GO terms enriched in this cluster were associated with chromosome segregation, mitotic nuclear division and cell cycle phase transition (Fig. 7D and Suppl. Table S8).

### DEGs-based transcription factor enrichment

To further investigate the potential upstream regulators of the abovementioned mechanisms, we performed a transcription factor (TF) binding site enrichment analysis based on the DEGs from our transcriptomic data. Specifically, we identified E2F family, NFYA, ESR1, MYC, SIN3A and TP53 among the significantly enriched TFs. We connected the enriched TFs to the irradiation-related gene sets predicted by GSEA in our dataset (Suppl. Table S9).

### Proteomics analysis revealed less pronounced effects of irradiation

To investigate whether the gene expression changes could be detectable on a protein level, we conducted a proteomic analysis using LC-MS/MS on both cPOCs (n=5) and mPOCs (n=5). PCA revealed that PC1 accounted for 44.34% and 41.02% of the variance in cPOCs and mPOCs, respectively. However, PC1 and PC2 did not differentiate between the irradiated time points and controls (Suppl. Fig. S6A, B).

We next identified differentially expressed peptides (DEPs) between the irradiated groups and their respective controls. As for transcriptomics analysis, DEPs were analysed across various time points: 1 h and 4 h post-irradiation compared to the 1 h control, and 24 h post-irradiation compared to the 24 h control. In cPOCs, irradiation had minimal effect at 1 h and 4 h with only 9 and 26 DEPs, respectively (Suppl. Fig. S6C). A more pronounced impact was observed at 24 h post-irradiation with 175 DEPs detected (Suppl. Fig. S6C). Similar to the transcriptomics data, we identified DEPs connected to cytoskeleton and ECM organisation (*i.e.*, TJP2, CNN1, COL3A1, COL4A2, COL2A1), metabolism (*i.e.*, ATP5F1E, GLUD1, ENO1) in cPOCs at different time points post-irradiation (Suppl. Table S10). In contrast, mPOCs showed 88 DEPs at 1 h, 21 at 4 h, and 139 DEPs at 24 h post-irradiation (Suppl. Fig. S6C). Consistent with our transcriptomics data, peptides related to cytoskeleton organisation (*i.e.*, ACTN1, CNN1, EPS8L2), oxygen and energy metabolism (*i.e.*, CYGB, ATP5F1E), and ECM (*i.e.*, COL23A1) were differentially expressed in mPOCs at different time points post-irradiation (Suppl. Table S10).

### Irradiation hinders the ability of cPOCs and mPOCs to aggregate *in vitro*

Although no significant cell death was observed throughout the recovery time points, we identified transcriptional changes in chromatin segregation, ECM and cytoskeleton organisation in both cPOCs and mPOCs. The alteration of these mechanisms suggests a hindered functionality in cell-cell interaction after X-ray exposure. To investigate this, we tested the ability of both cPOCs and mPOCs to form aggregates *in vitro*, by seeding cells onto Biosilk scaffolds (Di Nisio et al., 2024) 24 h post-irradiation. Non-irradiated cPOCs and mPOCs served as control group for comparison. The cells were cultured on the Biosilk scaffolds for 14 days, after which they were transferred to a floating culture system and observed for up to 40 days (Fig. 8A).

**Figure 8.**
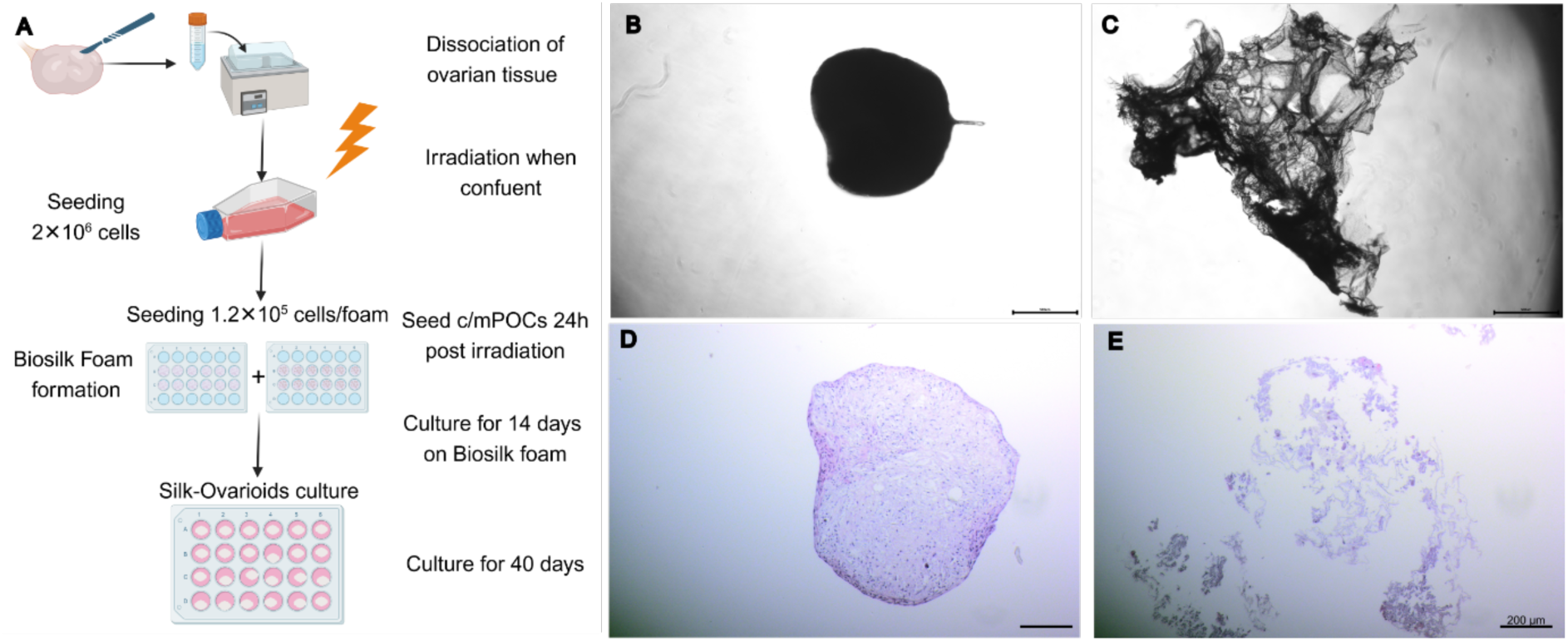
Functional *in vitro* assessment of capacity to aggregate in control and irradiated cPOCs and mPOCs: Silk-Ovarioids. **(A)** Schematic representation of the experimental approach for Silk-Ovarioids formation (n=2 patients; 6 BioSilk foams/patient). Graph generated using BioRender. Representative image of Silk-Ovarioids formation from **(B)** non-irradiated control POCs and **(C)** irradiated POCs after 40 days in free-floating culture. Scale bar = 500 μm. H&E staining of a 3D spheroid derived from **(D)** non-irradiated control POCs and **(E)** irradiated POCs. Scale bar = 200 μm.

The experiment was conducted using cPOCs and mPOCs derived from 2 different patients. For each patient, 6 foams were seeded to form Silk-Ovarioids in non-irradiated and irradiated groups. In the non-irradiated cPOCs group, all 12 foams resulted in Silk-Ovarioids formation (100%), while in the irradiated group, only 3 out of 12 foams produced Silk-Ovarioids (25%). Similarly, in the non-irradiated mPOCs group, 10 out of 12 foams led to Silk-Ovarioids formation (83%), whereas none of the irradiated mPOCs formed Silk-Ovarioids (0%) (Suppl. Table S11).

In the non-irradiated controls, well-defined Silk-Ovarioids were observed under an inverted phase-contrast microscope (Fig. 8B). Silk-Ovarioids formed within one and three days of free-floating culture in cPOCs and mPOCs controls, respectively. These structures remained stable and well-defined for up to 40 days (Fig. 8B). In contrast, most of the irradiated cPOCs and mPOCs, failed to form Silk-Ovarioids (Fig. 8C). Hematoxylin and Eosin (H&E) staining further confirmed these findings. In fact, Silk-Ovarioids from non-irradiated cPOCs and mPOCs appeared well-defined, containing intact cells within the aggregated structure (Fig. 8D). On the contrary, irradiated cPOCs and mPOCs were hardly visible on the silk fibers and, when present, displayed disorganised cellular architecture (Fig. 8E). Moreover, the Silk-Ovarioids from cPOCs that appeared intact initially, were disrupted during the process of fixation and staining.

## Discussion

This study aimed to identify for the first time the molecular signature of acute high-dose X-ray exposure on human primary ovarian cells isolated from both the cortex (cPOCs) and medulla (mPOCs). This novel approach contributes to our understanding of radiation-induced ovarian damage and provides the groundwork for future studies aimed at developing protective protocols for ovarian tissue in patients undergoing radiotherapy.

After screening different doses and time of recovery with KGN granulosa-like cancer cell line, we selected 10 Gy as the acute high-dose, still sublethal X-ray treatment. In the outlined experimental settings, we did not observe significant differences in ATP levels or mitochondrial activity in irradiated cPOCs and mPOCs. Intracellular ATP levels indicate cellular viability (Niles et al., 2021), while mitochondrial activity plays a critical role in regulating cellular proliferation (Antico Arciuch et al., 2012). Our findings suggest that a single 10 Gy dose of irradiation does not disrupt core cellular processes on the observed timescale, allowing cells to continue being viable and proliferating.

To assess if canonical pathways induced by irradiation were active in our experimental settings, we performed immunofluorescence staining on selected markers. We detected activation of DNA damage pathway at early time points after irradiation, as demonstrated by the upregulation of γ-H2AX and p-Chk1 in both cPOCs and mPOCs. In fact, literature data show that irradiation-induced DNA damage occurs through single- or double-strand breaks (SSBs or DSBs) or indirectly (∼70% of DNA damage) via free radicals generated during irradiation (Baskar et al., 2014; Wang et al., 2018). This activates the DNA damage response pathway, although approximately 5% of DSBs remain unrepaired (Minchom et al., 2018), leading to cell cycle arrest to prevent the replication of damaged DNA. As a response to DNA damage, γH2AX is activated at a very early stage (Siddiqui et al., 2015), and subsequently p-Chk1 initiates the mechanisms of DNA damage repair (Patil et al., 2013). In concordance with these DNA damage response mechanisms, at a transcriptomic level, we observed an early upregulation of p53 pathway, apoptosis, UV response up, and mTORC1 signaling at 4 h post-irradiation in both cPOCs and mPOCs. This can be further confirmed by upregulation of DEGs associated with p53-dependent apoptosis in cPOCs and mPOCs at 4 h according to the gene pattern analysis. In fact, the upregulation of p53 pathway is crucial for DNA damage response, leading to either cell cycle arrest or apoptosis (Reinhardt and Schumacher, 2012). In our experiments, we demonstrated initial increase of pro-apoptotic markers, namely the active forms of caspase 3 and p53, at the very early time point (1 h), followed by a steady reduction at the later culture times. This is supported by the overexpression of the anti-apoptotic marker Bcl-2 during the first 4 h post-irradiation, sustaining the hypothesis of the transition from apoptosis to cell cycle blockage. However, these results may be influenced by the unintended selection of surviving cells post-irradiation, a potential bias introduced by the technical procedures of immunostaining assays. Nevertheless, this p53-dependent mechanism has been previously described in immortalized primary retinal pigment epithelium cells, showing a slow recovery of mitosis after 10 Gy irradiation (Reyes et al., 2018).

In mice and human cell lines, it has been reported that radiosensitive tissues display a prolonged p53 signalling activation (Stewart-Ornstein et al., 2021). As well, this activation has been widely reported as a marker of radiation sensitivity in oncological patients (Stewart-Ornstein et al., 2021; Thariat et al., 2021). In our transcriptomic data, we detected a sustained upregulation of p53 pathway at 24 h post-irradiation, accompanied by overall repression of cell cycle related signals, such as E2F, mTORC1, G2M checkpoint and MYC targets. Consistent with the mechanism elucidated by Reyes et al. (Reyes et al., 2018) that outlined a p53-dependent cell cycle arrest through p-p21 activation and inhibition of cyclin E, we also identified the lack of p-p53 protein expression and the upregulation of p-p21 in both cPOCs and mPOCs. The mechanism that inhibits the expression of cyclin E seems to affect predominantly mPOCs, that display a significant downregulation of this selected cyclin. In addition, the steady downregulation of DEGs throughout the tested time points reveals a disruption of chromatin segregation and mitosis, as evidenced by the gene pattern analysis in cPOCs and mPOCs. Previous report suggests that effective repair of initial irradiation-induced DNA damage allows cells to continue progressing through the cell cycle. However, if repair is unsuccessful, cells may either undergo cell death and exit the cycle or persist while attempting to resolve genomic instabilities (Lonati et al., 2021). In addition, analysis at a protein level of cell cycle progression showed stable levels of cyclin D1 in cPOCs, while mPOCs exhibit decreasing levels of cyclin D1 up to 24 h post-irradiation. This suggests distinct cell cycle responses in cPOCs and mPOCs, further indicating that mPOCs are a more radiosensitive target.

One of the main contributors to cell cycle progression is the transcription factor Myc (García-Gutiérrez et al., 2019). Myc regulates cell cycle progression primarily by suppressing the expression or activity of key cell cycle inhibitors, such as p15, p21 and p27 (Gartel et al., 2001; Serrano et al., 1993). Therefore, the downregulation of Myc targets in our transcriptomic dataset, together with the upregulation of p-p21 may grant the cell cycle arrest mechanisms in both cPOCs and mPOCs. Furthermore, the G1 cell cycle arrest requires repression of Myc mediated by p53, through the recruitment of Sin3a, a Myc corepressor (Ho et al., 2005). Therefore, to better understand the putative role of Myc in our system, we performed the TF enrichment analyses using DEGs stratified by up- and downregulation identified at 24 h post-irradiation. Notably, MYC, SIN3A and TP53 were enriched in irradiation-induced upregulated genes of cPOCs and mPOCs. This observed molecular signature, together with the previously documented cell cycle arrest, suggests that the p53-mediated suppression of Myc’s transcription factor activity in cell cycle progression could serve as first-line response to X-ray exposure in primary ovarian cells. Particularly in mPOCs, this mechanism is corroborated by the early downregulation of gene patterns involving histone modification at 4 h post-irradiation, that accompanies the above mentioned Myc repression process (Ho et al., 2005). Even though Myc has been previously reported to induce p53-mediated apoptosis in mouse models (Phesse et al., 2014), our transcriptomic data displayed downregulation of *MYC* in the irradiated samples compared to their corresponding controls.

Through proteomics analysis we identified changes of ECM- and cytoskeleton-related peptides after irradiation. The ECM provides essential physical scaffolding and crucial signals for tissue morphogenesis, differentiation and homeostasis (Frantz et al., 2010). Integrins, as ECM receptors, mediate cell adhesion to the ECM (Harburger and Calderwood, 2009; Xian et al., 2010). Additionally, cell adhesion influences cytoskeleton formation and is involved in cell migration (Schmidt and Friedl, 2010). In this study, irradiation impaired the ability of cPOCs and mPOCs to form Silk-Ovarioids. Our previous study showed that Silk-Ovarioids derived from both cPOCs and mPOCs sustain long-term culture and their formation is strictly dependent on the successful cell-cell interaction and *de novo* ECM formation (Di Nisio et al., 2024). Overall, these findings indicate that irradiation significantly impacts ECM and cytoskeleton formation, leading to structural disorganisation, loss of cell adhesion and subsequent dysfunction in Silk-Ovarioid formation.

The present study expands our understanding on how irradiation impacts POCs derived from cortex and medulla, suggesting a Myc-p53 centered first-line response to acute high-dose X-ray exposure. Although this study utilised ovarian tissue from gender-affirmation patients who underwent androgen treatment prior to surgical removal, our previous study revealed a comparable ovarian cell composition with ovarian tissue retrieved from patients undergoing caesarean section (Wagner et al., 2020). However, potential differences due to the hormone treatment cannot be entirely excluded. Additionally, in this study the impact of X-ray exposure was assessed on a monolayer cell model, thus limiting the extrapolation power of our results to ovary *in vivo*. Lastly, the impact of irradiation was focused on the somatic cell populations that are essential for ovarian function. Further studies are needed to investigate the effects of X-ray exposure on ovarian follicles and their function.

In conclusion, this study provides novel insights into the response of cPOCs and mPOCs to irradiation. We demonstrate that a single 10 Gy irradiation dose is sufficient to induce sublethal damage, with main effects on cell cycle arrest, cell adhesion and interaction. Our findings suggest altered molecular signature of first-line response to irradiation in human primary ovarian cells. These findings are crucial for understanding radiation-induced ovarian damage. This paves the way for future clinically relevant studies that may focus on MYC-p53 pathways and interactions in response to irradiation and their potential use towards the identification of ovarian “fertoprotective” strategies during radiotherapy.

## Supplementary Tables and table legends

The data are provided in the submitted Excel files.

**Supplementary Table S1:** Seeding density information for each cell type, categorized by experimental approach.

**Supplementary Table S2:** Overview of primary and secondary antibodies used for immunofluorescence staining.

**Supplementary Table S3:** Information on upregulated and downregulated DEGs in (**Sheet 1**) 24 h control cPOCs compared to 1 h control. (**Sheet 2**) 24 h control mPOCs compared to 1 h control.

**Supplementary Table S4:** Information on upregulated and downregulated DEGs in (**Sheet 1**) Irradiated (IR) 1 h cPOCs compared to 1 h Control cPOCs. (**Sheet 2**) IR 4 h cPOCs compared to 1 h Control cPOCs. (**Sheet 3**) IR 24 h cPOCs compared to 24 h Control cPOCs. (**Sheet 4**) IR 1 h mPOCs compared to 1 h Control mPOCs. (**Sheet 5**) IR 4 h mPOCs compared to 1 h Control mPOCs. (**Sheet 6**) IR 24 h mPOCs compared to 24 h Control mPOCs.

**Supplementary Table S5**: Information of Gene Set Enrichment Analysis (GSEA) in (**Sheet 1**) 24 h control cPOCs compared to 1 h control. (**Sheet 2**) 24 h control mPOCs compared to 1 h control.

**Supplementary Table S6**: Information of Gene Set Enrichment Analysis (GSEA) of all expressed genes in (**Sheet 1**) Irradiated (IR) 1 h cPOCs compared to 1 h control cPOCs. (**Sheet 2**) IR 4 h cPOCs compared to 1 h control cPOCs. (**Sheet 3**) IR 24 h cPOCs compared to 24 h control cPOCs. (**Sheet 4**) IR 1 h mPOCs compared to 1 h control mPOCs. (**Sheet 5**) IR 4 h mPOCs compared to 1 h control mPOCs. (**Sheet 6**) IR 24 h mPOCs compared to 24 h control mPOCs.

**Supplementary Table S7:** Information on the gene pattern enriched GO against all annotated genes for (**Sheet 1**) cluster 1 in cPOCs. (**Sheet 2**) cluster 2 in cPOCs. (**Sheet 3**) cluster 3 in cPOCs. (**Sheet 4**) cluster 4 in cPOCs. (**Sheet 5**) cluster 5 in cPOCs. (**Sheet 6**) cluster 6 in cPOCs.

**Supplementary Table S8:** Information on the gene pattern enriched GO against all annotated genes for (**Sheet 1**) cluster 1 in mPOCs. (**Sheet 2**) cluster 2 in mPOCs. (**Sheet 3**) cluster 3 in mPOCs. (**Sheet 4**) cluster 4 in mPOCs. (**Sheet 5**) cluster 5 in mPOCs. (**Sheet 6**) cluster 6 in mPOCs.

**Supplementary Table S9:** Information on (**Sheet 1**) downregulated transcription factors (TF) in cPOCs 24 h post-irradiation. (**Sheet 2**) downregulated TF overlap with motifs in cPOCs 24 h post-irradiation. (**Sheet 3**) upregulated TF in cPOCs 24 h post-irradiation. (**Sheet 4**) upregulated TF overlap with motifs in cPOCs 24 h post-irradiation. (**Sheet 5**) downregulated TF in mPOCs 24 h post-irradiation. (**Sheet 6**) downregulated TF overlap with motifs in mPOCs 24 h post-irradiation. (**Sheet 7**) upregulated TF in mPOCs 24 h post-irradiation. (**Sheet 8**) upregulated TF overlap with motifs in mPOCs 24 h post-irradiation. (**Sheet 9**) up- and downregulated TF and motifs in cPOCs 24 h post-irradiation and their connection in GSEA from our dataset. (**Sheet 10**) up- and downregulated TF and motifs in mPOCs 24 h post-irradiation and their connection in GSEA from our dataset.

**Supplementary Table S10:** Information on upregulated and downregulated DEPs at (**Sheet 1**) Irradiated (IR) 1 h cPOCs compared to 1 h Control cPOCs. (**Sheet 2**) IR 4 h cPOCs compared to 1 h Control cPOCs. (**Sheet 3**) IR 24 h cPOCs compared to 24 h Control cPOCs. (**Sheet 4**) IR 1 h mPOCs compared to 1 h Control mPOCs. (**Sheet 5**) IR 4 h mPOCs compared to 1 h Control mPOCs. (**Sheet 6**) IR 24 h mPOCs compared to 24 h Control mPOCs.

**Supplementary Table S11**: Overview of patients, number of cells and days of culture required for the 3D Silk-Ovarioids Formation.

## Ethical Approval

The use of ovarian tissue in research was approved by the Stockholm Region Ethical Review Board (license number 2024-08606-01). Clinicians informed the patients about the study, and ovarian tissue was biopsied from ovaries collected from patients undergoing gender-affirming surgeries after written and signed informed consent.

## Consent to Publish

All the authors have approved the manuscript and agree to its publication.

## Authors Contribution

SPD, AS and VDN contributed to the study’s conception. SPD, DL, AVM, PD, AS and VDN contributed to the experimental design. SPD and VDN performed the laboratory work of the study. SPD, TL and VDN performed the visualisation of the data used in this study. AV and RZ performed the proteomics. SPD collected the data. SPD, EML and TL performed the statistical analysis of the data. TL and AD performed the RNA-sequencing and proteomics analysis. KP coordinated patient recruitment and evaluated patients’ health. GA, PD and AS were responsible of supervision and funding acquisition. SPD wrote the first draft. SPD, TL and VDN wrote the final version of the manuscript. All the authors read, commented and approved the final version of the manuscript.

## Funding

This work was funded by the European Union’s HORIZON 2020 research and innovation programme (MATER) under the Marie Skłodowska-Curie Actions (grant agreement No: 813707), the Estonian Research Council (grants PRG1076 and PSG608), the Orion Research Foundation sr personal grant, the Research grant from the Center for Innovative Medicine (CIMED) and the Consolidator Grant at Karolinska Institutet.

## Conflict of Interest

The authors declare no competing interests.

## Availability of data

RNA-sequencing count matrix is deposited in Gene Expression Omnibus (GEO) with accession number GSE291604. The mass spectrometry proteomics data have been deposited to the ProteomeXchange Consortium via the PRIDE (Perez-Riverol et al, 2022) partner repository with the dataset identifier PXD061796. The code used for the analysis can be found in https://github.com/tialiv/X-Ovary.

## Acknowledgments

The authors would like to express their gratitude to the clinicians and research nurses who supported patient recruitment and biopsy collection, as well as to the patients whose invaluable contribution made this research possible. This study was partially performed at the Live Cell Imaging Core facility/Nikon Center of Excellence, at Karolinska Institutet, Sweden, supported by the KI infrastructure council. We would like to thank BEA, the Bioinformatics and Expression Analysis core facility, and FENO which is supported by the board of research at the Karolinska Institutet. The authors acknowledge support from the National Genomics Infrastructure in Stockholm funded by Science for Life Laboratory, the Knut and Alice Wallenberg Foundation and the Swedish Research Council, and SNIC/NAISS/Uppsala Multidisciplinary Center for Advanced Computational Science for assistance with massively parallel sequencing and access to the UPPMAX computational infrastructure. Protein identification and quantification were carried out by the Proteomics Biomedicum core facility, Karolinska Institutet (https://ki.se/en/research/proteomics-biomedicum-core-facility). We wish to thank the Biobank and Study Support at Karolinska University Hospital for their contribution including professional service and support. The authors would like to thank the Orion Research Foundation sr for providing the personal grant for the first author to conduct this study.

## Notes

### Competing Interest Statement

The authors have declared no competing interest.

## References

Antico Arciuch, V.G., Elguero, M.E., Poderoso, J.J., Carreras, M.C., 2012. Mitochondrial Regulation of Cell Cycle and Proliferation. Antioxid Redox Signal 16, 1150–1180. 10.1089/ars.2011.4085

Barton, M.B., Jacob, S., Shafiq, J., Wong, K., Thompson, S.R., Hanna, T.P., Delaney, G.P., 2014. Estimating the demand for radiotherapy from the evidence: A review of changes from 2003 to 2012. Radiotherapy and Oncology 112, 140–144. 10.1016/j.radonc.2014.03.024

Baskar, R., Dai, J., Wenlong, N., Yeo, R., Yeoh, K.-W., 2014. Biological response of cancer cells to radiation treatment. Front. Mol. Biosci. 1. 10.3389/fmolb.2014.00024

Bentzen, S.M., Heeren, G., Cottier, B., Slotman, B., Glimelius, B., Lievens, Y., van den Bogaert, W., 2005. Towards evidence-based guidelines for radiotherapy infrastructure and staffing needs in Europe: the ESTRO QUARTS project. Radiother Oncol 75, 355–365. 10.1016/j.radonc.2004.12.007

Cao, W., Qin, K., Li, F., Chen, W., 2024. Comparative study of cancer profiles between 2020 and 2022 using global cancer statistics (GLOBOCAN). J Natl Cancer Cent 4, 128–134. 10.1016/j.jncc.2024.05.001

Chemaitilly, W., Mertens, A.C., Mitby, P., Whitton, J., Stovall, M., Yasui, Y., Robison, L.L., Sklar, C.A., 2006. Acute Ovarian Failure in the Childhood Cancer Survivor Study. The Journal of Clinical Endocrinology & Metabolism 91, 1723–1728. 10.1210/jc.2006-0020

Fan, X., Bialecka, M., Moustakas, I., Lam, E., Torrens-Juaneda, V., Borggreven, N.V., Trouw, L., Louwe, L.A., Pilgram, G.S.K., Mei, H., Westerlaken, L. van der, Lopes, S.M.C. de S., 2019. Single-cell reconstruction of follicular remodeling in the human adult ovary. Nature Communications 10, 3164. 10.1038/s41467-019-11036-9

Fernström, E., Jarfelt, M., Blomstrand, M., Lannering, B., Axelsson, M., Wasling, P., Björk-Eriksson, T., Zetterberg, H., Kalm, M., 2024. CSF biomarkers of neurotoxicity in childhood cancer survivors after cranial radiotherapy or surgery. Annals of Clinical and Translational Neurology 11, 2382. 10.1002/acn3.52152

Frantz, C., Stewart, K.M., Weaver, V.M., 2010. The extracellular matrix at a glance. J Cell Sci 123, 4195–4200. 10.1242/jcs.023820

García-Gutiérrez, L., Delgado, M.D., León, J., 2019. MYC Oncogene Contributions to Release of Cell Cycle Brakes. Genes 10, 244. 10.3390/genes10030244

Gartel, A.L., Ye, X., Goufman, E., Shianov, P., Hay, N., Najmabadi, F., Tyner, A.L., 2001. Myc represses the p21(WAF1/CIP1) promoter and interacts with Sp1/Sp3. Proceedings of the National Academy of Sciences 98, 4510–4515. 10.1073/pnas.081074898

Gu, Z., 2022. Complex heatmap visualization. Imeta 1, e43. 10.1002/imt2.43

Gu, Z., Eils, R., Schlesner, M., 2016. Complex heatmaps reveal patterns and correlations in multidimensional genomic data. Bioinformatics 32, 2847–2849. 10.1093/bioinformatics/btw313

Harburger, D.S., Calderwood, D.A., 2009. Integrin signalling at a glance. J Cell Sci 122, 159–163. 10.1242/jcs.018093

Ho, J.S.L., Ma, W., Mao, D.Y.L., Benchimol, S., 2005. p53-Dependent Transcriptional Repression of c-myc Is Required for G1 Cell Cycle Arrest. Mol Cell Biol 25, 7423–7431. 10.1128/MCB.25.17.7423-7431.2005

Immediata, V., Ronchetti, C., Spadaro, D., Cirillo, F., Levi-Setti, P.E., 2022. Oxidative Stress and Human Ovarian Response—From Somatic Ovarian Cells to Oocytes Damage: A Clinical Comprehensive Narrative Review. Antioxidants 11, 1335. 10.3390/antiox11071335

Irtan, S., Orbach, D., Helfre, S., Sarnacki, S., 2013. Ovarian transposition in prepubescent and adolescent girls with cancer. The Lancet Oncology 14, e601–e608. 10.1016/S1470-2045(13)70288-2

Kimler, B.F., Briley, S.M., Johnson, B.W., Armstrong, A.G., Jasti, S., Duncan, F.E., 2018. Radiation-induced ovarian follicle loss occurs without overt stromal changes. Reproduction 155, 553–562. 10.1530/REP-18-0089

Lawrie, T.A., Evans, J., Gillespie, D., Erridge, S., Vale, L., Kernohan, A., Grant, R., 2018. Long-term side effects of radiotherapy, with or without chemotherapy, for glioma. The Cochrane Database of Systematic Reviews 2018, CD013047. 10.1002/14651858.CD013047

Liberzon, A., Subramanian, A., Pinchback, R., Thorvaldsdóttir, H., Tamayo, P., Mesirov, J.P., 2011. Molecular signatures database (MSigDB) 3.0. Bioinformatics 27, 1739–1740. 10.1093/bioinformatics/btr260

Lonati, L., Barbieri, S., Guardamagna, I., Ottolenghi, A., Baiocco, G., 2021. Radiation-induced cell cycle perturbations: a computational tool validated with flow-cytometry data. Sci Rep 11, 925. 10.1038/s41598-020-79934-3

Luo, W., Brouwer, C., 2013. Pathview: an R/Bioconductor package for pathway-based data integration and visualization. Bioinformatics 29, 1830–1831. 10.1093/bioinformatics/btt285

Meirow, D., Nugent, D., 2001. The effects of radiotherapy and chemotherapy on female reproduction. Human Reproduction Update 7, 535–543. 10.1093/humupd/7.6.535

Minchom, A., Aversa, C., Lopez, J., 2018. Dancing with the DNA damage response: next-generation anti-cancer therapeutic strategies. Ther Adv Med Oncol 10, 1758835918786658. 10.1177/1758835918786658

Mishra, B., Ripperdan, R., Ortiz, L., Luderer, U., 2017. Very low doses of heavy oxygen ion radiation induce premature ovarian failure. Reproduction 154, 123–133. 10.1530/REP-17-0101

Mulder, R.L., Font-Gonzalez, A., Hudson, M.M., van Santen, H.M., Loeffen, E.A.H., Burns, K.C., Quinn, G.P., van Dulmen-den Broeder, E., Byrne, J., Haupt, R., Wallace, W.H., van den Heuvel-Eibrink, M.M., Anazodo, A., Anderson, R.A., Barnbrock, A., Beck, J.D., Bos, A.M.E., Demeestere, I., Denzer, C., Di Iorgi, N., Hoefgen, H.R., Kebudi, R., Lambalk, C., Langer, T., Meacham, L.R., Rodriguez-Wallberg, K., Stern, C., Stutz-Grunder, E., van Dorp, W., Veening, M., Veldkamp, S., van der Meulen, E., Constine, L.S., Kenney, L.B., van de Wetering, M.D., Kremer, L.C.M., Levine, J., Tissing, W.J.E., Berger, C., Diesch, T., Dirksen, U., Ginsberg, J., Giwercman, A., Grabow, D., Gracia, C., Hunter, S.E., Inthorn, J., Kaatsch, P., Kelvin, J.F., Klosky, J.L., Laven, J.S.E., Lockart, B.A., Neggers, S.J., Paul, N.W., Peate, M., Phillips, B., Reed, D.R., Tinner, E.M.E., van den Berg, M., Verhaak, C., 2021. Fertility preservation for female patients with childhood, adolescent, and young adult cancer: recommendations from the PanCareLIFE Consortium and the International Late Effects of Childhood Cancer Guideline Harmonization Group. The Lancet Oncology 22, e45–e56. 10.1016/S1470-2045(20)30594-5

Niles, A.L., Cali, J.J., Lazar, D.F., 2021. A live-cell assay for the real-time assessment of extracellular ATP levels. Analytical Biochemistry 628, 114286. 10.1016/j.ab.2021.114286

Nisio, V.D., Li, T., Xiao, Z., Papaikonomou, K., Damdimopoulos, A., Végvári, Á., Lebre, F., Alfaro-Moreno, E., Pedersen, M., Svingen, T., Zubarev, R., Acharya, G., Damdimopoulou, P., Salumets, A., 2024. Silk-Ovarioids: Establishment and characterization of human ovarian primary cells 3D-model system. 10.1101/2024.07.31.606024

Pantano, L., 2024. DEGreport [WWW Document]. Bioconductor. URL http://bioconductor.org/packages/DEGreport/ (accessed 1.16.25).

Patil, M., Pabla, N., Dong, Z., 2013. Checkpoint kinase 1 in DNA damage response and cell cycle regulation. Cell Mol Life Sci 70, 4009–4021. 10.1007/s00018-013-1307-3

Phesse, T.J., Myant, K.B., Cole, A.M., Ridgway, R.A., Pearson, H., Muncan, V., van den Brink, G.R., Vousden, K.H., Sears, R., Vassilev, L.T., Clarke, A.R., Sansom, O.J., 2014. Endogenous c-Myc is essential for p53-induced apoptosis in response to DNA damage in vivo. Cell Death Differ 21, 956–966. 10.1038/cdd.2014.15

Pouvreau, P., Taleb, I., Fontaine, A., Edouard, L., Gibson, N., Yaouanq, M., Boudoussier, A., Petit, A., Vinh-Hung, V., Sargos, P., Benziane-Ouaritini, N., Bouleftour, W., Magne, N., 2024. Heart is a heavy burden: cardiac toxicity in radiation oncology. Support Care Cancer 32, 769. 10.1007/s00520-024-08949-7

R Core Team, 2022. R: A Language and Environment for Statistical Computing | BibSonomy [WWW Document]. https://www.R-project.org/. URL https://www.bibsonomy.org/bibtex/7469ffee3b07f9167cf47e7555041ee7 (accessed 1.16.25).

Reinhardt, H.C., Schumacher, B., 2012. The p53 network: Cellular and systemic DNA damage responses in aging and cancer. Trends Genet 28, 128–136. 10.1016/j.tig.2011.12.002

Reyes, J., Chen, J.-Y., Stewart-Ornstein, J., Karhohs, K.W., Mock, C.S., Lahav, G., 2018. Fluctuations in p53 Signaling Allow Escape from Cell-Cycle Arrest. Molecular Cell 71, 581–591.e5. 10.1016/j.molcel.2018.06.031

Ritchie, M.E., Phipson, B., Wu, D., Hu, Y., Law, C.W., Shi, W., Smyth, G.K., 2015. limma powers differential expression analyses for RNA-sequencing and microarray studies. Nucleic Acids Research 43, e47. 10.1093/nar/gkv007

Roos, K., Rooda, I., Keif, R.-S., Liivrand, M., Smolander, O.-P., Salumets, A., Velthut-Meikas, A., 2022. Single-cell RNA-seq analysis and cell-cluster deconvolution of the human preovulatory follicular fluid cells provide insights into the pathophysiology of ovarian hyporesponse. Front Endocrinol (Lausanne) 13, 945347. 10.3389/fendo.2022.945347

Schmidt, K.L.T., Ernst, E., Byskov, A.G., Nyboe Andersen, A., Yding Andersen, C., 2003. Survival of primordial follicles following prolonged transportation of ovarian tissue prior to cryopreservation. Human Reproduction 18, 2654–2659. 10.1093/humrep/deg500

Schmidt, S., Friedl, P., 2010. Interstitial cell migration: integrin-dependent and alternative adhesion mechanisms. Cell Tissue Res 339, 83–92. 10.1007/s00441-009-0892-9

Serrano, M., Hannon, G.J., Beach, D., 1993. A new regulatory motif in cell-cycle control causing specific inhibition of cyclin D/CDK4. Nature 366, 704–707. 10.1038/366704a0

Shin, E., Lee, S., Kang, H., Kim, J., Kim, K., Youn, H., Jin, Y.W., Seo, S., Youn, B., 2020. Organ-Specific Effects of Low Dose Radiation Exposure: A Comprehensive Review. Front Genet 11, 566244. 10.3389/fgene.2020.566244

Siddiqui, M.S., François, M., Fenech, M.F., Leifert, W.R., 2015. Persistent γH2AX: A promising molecular marker of DNA damage and aging. Mutation Research/Reviews in Mutation Research 766, 1–19. 10.1016/j.mrrev.2015.07.001

Stewart-Ornstein, J., Iwamoto, Y., Miller, M.A., Prytyskach, M.A., Ferretti, S., Holzer, P., Kallen, J., Furet, P., Jambhekar, A., Forrester, W.C., Weissleder, R., Lahav, G., 2021. p53 dynamics vary between tissues and are linked with radiation sensitivity. Nat Commun 12, 898. 10.1038/s41467-021-21145-z

Stirling, D.R., Swain-Bowden, M.J., Lucas, A.M., Carpenter, A.E., Cimini, B.A., Goodman, A., 2021. CellProfiler 4: improvements in speed, utility and usability. BMC Bioinformatics 22, 433. 10.1186/s12859-021-04344-9

Stroud, J.S., Mutch, D., Rader, J., Powell, M., Thaker, P.H., Grigsby, P.W., 2009. Effects of cancer treatment on ovarian function. Fertility and Sterility 92, 417–427. 10.1016/j.fertnstert.2008.07.1714

Tarvainen, I., Soto, D.A., Laws, M.J., Björvang, R.D., Damdimopoulos, A., Roos, K., Li, T., Kramer, S., Li, Z., Lavogina, D., Visser, N., Kallak, T.K., Lager, S., Gidlöf, S., Edlund, E., Papaikonomou, K., Öberg, M., Olovsson, M., Salumets, A., Velthut-Meikas, A., Flaws, J.A., Damdimopoulou, P., 2023. Identification of phthalate mixture exposure targets in the human and mouse ovary in vitro. Reprod Toxicol 119, 108393. 10.1016/j.reprotox.2023.108393

Thariat, J., Chevalier, F., Orbach, D., Ollivier, L., Marcy, P.-Y., Corradini, N., Beddok, A., Foray, N., Bougeard, G., 2021. Avoidance or adaptation of radiotherapy in patients with cancer with Li-Fraumeni and heritable TP53-related cancer syndromes. The Lancet Oncology 22, e562–e574. 10.1016/S1470-2045(21)00425-3

Wagner, M., Yoshihara, M., Douagi, I., Damdimopoulos, A., Panula, S., Petropoulos, S., Lu, H., Pettersson, K., Palm, K., Katayama, S., Hovatta, O., Kere, J., Lanner, F., Damdimopoulou, P., 2020. Single-cell analysis of human ovarian cortex identifies distinct cell populations but no oogonial stem cells. Nat Commun 11, 1147. 10.1038/s41467-020-14936-3

Wallace, W.H.B., Thomson, A.B., Kelsey, T.W., 2003. The radiosensitivity of the human oocyte. Human Reproduction 18, 117–121. 10.1093/humrep/deg016

Wang, J., Wang, H., Qian, H., 2018. Biological effects of radiation on cancer cells. Military Medical Research 5, 20. 10.1186/s40779-018-0167-4

Wang, Y., Jiang, J., Zhang, J., Fan, P., Xu, J., 2023. Research Progress on the Etiology and Treatment of Premature Ovarian Insufficiency. Biomed Hub 8, 97–107. 10.1159/000535508

Wu, T., Hu, E., Xu, S., Chen, M., Guo, P., Dai, Z., Feng, T., Zhou, L., Tang, W., Zhan, L., Fu, X., Liu, S., Bo, X., Yu, G., 2021. clusterProfiler 4.0: A universal enrichment tool for interpreting omics data. Innovation (Camb) 2, 100141. 10.1016/j.xinn.2021.100141

Xian, X., Gopal, S., Couchman, J.R., 2010. Syndecans as receptors and organizers of the extracellular matrix | Cell and Tissue Research [WWW Document]. URL https://link.springer.com/article/10.1007/s00441-009-0829-3 (accessed 2.24.25).

Yu, G., Wang, L.-G., Han, Y., He, Q.-Y., 2012. clusterProfiler: an R package for comparing biological themes among gene clusters. OMICS 16, 284–287. 10.1089/omi.2011.0118

Yu, G., Wang, L.-G., Yan, G.-R., He, Q.-Y., 2015. DOSE: an R/Bioconductor package for disease ontology semantic and enrichment analysis. Bioinformatics 31, 608–609. 10.1093/bioinformatics/btu684

Zhu, A., Ibrahim, J.G., Love, M.I., 2019. Heavy-tailed prior distributions for sequence count data: removing the noise and preserving large differences. Bioinformatics 35, 2084–2092. 10.1093/bioinformatics/bty895

